# Chromosome-level genome assembly provides insights into adaptive evolution of chromosome and important traits in the gecko *Gekko japonicus*

**DOI:** 10.1101/2023.04.01.535199

**Authors:** Yinwei Wang, Youxia Yue, Chao Li, Zhiyi Chen, Yao Cai, Chaochao Hu, Yanfu Qu, Hong Li, Kaiya Zhou, Jie Yan, Peng Li

**Author notes:** These authors contributed equally to this work (Yinwei Wang, Youxia Yue). **Corresponding author: Dr. Jie Yan**, College of Life Sciences, Nanjing Normal University, No.1 Wenyuan Road, Nanjing, Jiangsu, 210023, P. R. China; **Dr. Peng Li**, College of Life Sciences, Nanjing Normal University, No.1 Wenyuan Road, Nanjing, Jiangsu, 210023, P. R. China. Competing Interests: The authors declared that no competing interests exist.

## Abstract

*Gekko japonicus* possess excellent flexible climbing and detoxification ability under insectivorous habits, and its chromosomes and the genetic evolutionary mechanisms behind these traits are still unclarified. Here, we assembled a chromosome-level genome of *G. japonicus* with a total size of 2.53 Gb contained in 19 pairs of chromosomes. The evolutionary breakpoint regions (EBRs) are significantly enriched for some repetitive elements compared to the rest of genome and the genes located in the EBRs are enriched in defense response pathway. *G. japonicus* specific gene families, expanded gene families and positively selected genes are mainly enriched in some pathways related to the immune, sensory and nervous systems. These results from comprehensive comparative genomics and evolutionary genomics analyses indicated that bitter taste receptor type 2 (T2Rs) expanded in different lineages by tandem gene duplication. The expansion and independent duplication events of T2Rs and positively selected branches were predominantly present in insectivorous species, suggests that T2Rs are associate with clearance of bitter toxins in gekkotans. Detoxification genes in detox and biosynthetic cytochrome P450 of *G. japonicas* have frequent duplication and loss events, suggests that they undergo more birth and death processes compared to biosynthesis type genes. Pro, Cys, Gly and Ser are the most abundant amino acids in 66 epidermis formation corneous beta proteins (CBPs) of *G. japonicas*, the abundance of Gly and Cys in CBPs implying significant effects on the flexibility and setae adhesiveness of gekkotans. Some thermosensitive thermoregulatory transient receptor potential channels under relaxed purifying selection or positive selection in *G. japonicus*, implying that one of the important factors improve the ability to adapt to climate change.

## Introduction

*Gekko japonicus*, as a member of the Gekkoniade which has over 1000 species, is widely distributed in Aisa (including China, Japan and Korea) (Toda and Yoshida 2005; Luu et al. 2017; Kim et al. 2019). Chromosome evolution plays important role on speciation. Studies on the chromosomes of Geckoidae have mainly focused on karyotype analysis, mainly including some standard Giemsa staining experiments, etc. It has been shown that the karyotype of most species of the genus *Gecko* is now found to be 2n = 38 (Shibaike et al. 2009; Qin et al. 2012). However, it is not enough to study chromosomes based on experiments alone, a series of bioinformatics-based analyses are also necessary. We assembled the genome of *G. japonicus* at the chromosome level based on Illumina Hiseq, Pacbio SMRT sequencing and Hi-C technologies, and the chromosome numbers obtained were consistent with previous experimental results (Shibaike et al. 2009). Our genome assembly quality and gene annotation are greatly improved compared to previous version and are well suited for chromosome level analysis. Due to the limited chromosome-level genome assembly of reptiles and the scarcity of those with high-quality gene annotations, we selected two other sauropsids, *Anolis carolinensis* and *Gallus gallus*, with relatively good assembly quality, complete gene annotations, and some variation in chromosome numbers, to compare with *G. japonicus* for synteny alignment and a series of analyses about evolutionary breakpoint regions (EBRs), which are the regions between two neighboring homologous synteny blocks (HSBs) and are considered to be species- or lineage-specific chromosome evolution related regions. EBR typically have higher GC content, repetitive elements and gene density compared to genome, although this is not always the case. The genes enriched in EBRs and the associated pathways and repetitive elements are thought to be important for chromosome evolution (Fan et al. 2019). Enrichment analyses of different gene sets in *G. japonicus*, including species-specific genes, species-specific expansion genes, species-specific positively selected genes, and species-specific EBRs, all revealed the evolution of specific adaptations in its immune, sensory, and nervous systems. This may be the underlying reason why *G. japonicus* is able to be nocturnal, hunt for prey through different senses, feed, escape from natural enemies and regenerate its tail etc.

Taste receptors, which are important for feeding in *G. japponicus*, are divided into two subfamilies, taste receptor type 1 (T1R) and taste receptor type 2 (T2R), and are widely found in vertebrates (Bachmanov and Beauchamp 2007). T1R is mainly involved in the perception of sweet and umami tastes, and its number varies little and is very conserved in reptiles (Feng and Liang 2018), whereas T2R is mainly involved in the perception of bitter tastes, and its number varies significantly throughout vertebrates compared to T1R, as evidenced by recent studies on the identification of bitter taste receptors in vertebrates and squamates (Li and Zhang 2014; Zhong et al. 2019). The Schlegel’s Japanese gecko *G. japonicus*, which primarily preys on insects, has a significant duplication and expansion of T2R genes, and in the enrichment analysis in this study, the genes enriched into taste-related pathways are mainly T2R. The location of T2R on the chromosome of *G. japonicus*, and the selection pressure on it are still unclear. In this study, the gene repertoire of *G. japonicus* was re-identified as well as evolutionary analysis was performed to reveal the duplication pattern and the selection pressure in different regions of T2R in *G. japonicus*.

Cytochrome P450 (CYP) is a very large gene superfamily that is widely distributed among all groups of organisms, and members from this superfamily are monooxygenases that act on a wide variety of substrates (Danielson 2002; Shankar and Mehendale 2014; Nelson 2018). In order to have a better classification and nomenclature for CYP, “clan” is defined as a higher-order category of the CYP family, and 11 clans were found in metazoans, including CYP clans 2, 3, 4, mitochondrial, 7, 19, 20, 26, 46, 51 and 74 (Nelson et al. 2013). Howerver, only ten clans were found in human and clan 74 has not beed found in vertebrates (Kawashima and Satta 2014). In function, CYPs are mainly involved in the synthesis and decomposition of a series of compounds, e.g. the synthesis of hormones and the decomposition of toxic substances, and are therefore mainly divided into two categories: biosynthesis type (B-type) and detoxification type (D-type). Four CYP gene families, CYP1, CYP2, CYP3, and CYP4, are considered D-type, while the others are B-type (Nebert and Dalton 2006). Studies on CYPs of vertebrate and invertebrate have shown that D-type CYP genes exhibit higher evolutionary instability and have more frequent birth and death processes compared to D-type CYP genes (Thomas 2007; Darragh et al. 2021). In birth and death model, the role that natural selection plays in the birth and death process is very complex; for example, in some taxa, detoxification genes were widely expanded and many positive selection signals were found, however, in some taxa, this was not the case (Dermauw et al. 2020). The gene repertoire, birth and death processes and evolutionary differences between D-type and B-type genes of CYPs in sauropsids have not been studied.

Like various other geckos, *G. japonicus* has some interesting traits such as small body size, agility and nocturnal habits, some of which have been mentioned above. Genes associated with some of these traits, which have been analyzed in previous study based on some candidate genes, including beta-keratin, opsin and olfactory receptor genes (Liu et al. 2015a). The study firstly focus on the corneous beta proteins (CBPs) family. Adaptive evolution of the amniotes from aquatic to terrestrial ecosystems has made them diverse in morphology, such as feathers of Aves, scales of Squamata, and shells of Testudines. In addition, epidermis is one of the important phenotypes in morphological identification, the evolution of which is one of great significance for amniotes to prevent mechanical damage and water loss. Epidermis and its appendages, e.g. feathers, claws, scales, etc., are composed of corneous beta proteins and some other relative proteins, the former of which used to be known as “β-keratins”. The β-keratins belong to the keratins which are the members of the intermediate filament protein family (Calvaresi et al. 2016) while CBPs represent the more abundant class of non-keratin proteins of sauropsids that associate to keratins as “keratin-associated beta-proteins” (KABPs) (Alibardi 2015). When it comes to keratinization and/orcornification, CBPs have been used more frequently in recent researches to distinguish them between the “true” keratins. The CBPs are encoded by a cluster of genes which are located between the genes for Loricrin (Lor) and Cornulin (Cn) in the centre of the epidermal differentiation complex (EDC) region in the chromosome, whereas the genes of S100A family are localized at the both borders of the EDC region. In fact, in addition to the genes mentioned above, numerous of other genes that are related to epidermal differentiation are located in EDC region as well. The sequences of CBP genes are conserved and each gene commonly contains two exons and one intron. The coding region derives from part of the second exon in 3’-terminal and produces a beta-protein containing five amino acid regions which are N-region, PRB (or precore box) region, CB (or core box) region, PC (or post-core box) region, and C-region (Alibardi 2016). It is reported that the extremely conserved core box sequence is the characteristic of CBPs. The expansion of CBPs in *G. japonicus* has been confirmed, and here, we re-analyzed CBPs, including gene re-identification, amino acid composition, gene conserved structure, phylogenetic and synteny analyses to supplement previous studies on CBPs in *G. japonicus* and other sauropsids (Liu et al. 2015a).

Body temperature is one of the most important variables affecting activities, foraging, enemy avoidance and reproduction of ectotherms. Ectotherms regulate body temperature under natural conditions through a variety of behavioral and physiological activities to maintain them in a certain temperature range (Angilletta et al. 1999). As a poikilothermal and nocturnal gecko, thermoregulation has been shown to play an important role for *Gekko japonicus*. Seasonal, diurnal temperature changes regulate the activity of gekkotans (Hitchcock and McBrayer 2006; Kim et al. 2018). However, up to date the molecular mechanisms of thermoregulation and related genes in gekkotans remain a mystery. Transient receptor potential (TRP) channels are a class of protein superfamily with six transmembrane segments that are widely distributed in green algae, fungi, choanoflagellates and a number of other eukaryotes and play important roles in a variety of physiological activities, such as lysosomal function, inflammatory response and thermoregulation (Venkatachalam and Montell 2007; Nilius and Owsianik 2011; Himmel and Cox 2020). The paucity of gene repertoire composition, phylogeny and evolutionary analyses of TRPs among sauropsids is largely attributed to the lack of genomes and the poor quality of assembly and annotation. In a recent study, the TRP gene repertoire composition, gene duplication events, and differential retention in tuatara were revealed for a subset of sauropsids, and a signature of positive selection in heat-sensitive genes TRPM2 and TRPV2 in tuatara was identified (Gemmell et al. 2020). Here, we increased more genomic data of sauropsids species to identify TRP genes and perform phylogenetic and evolutionary analyses to correct some of the results of previous study (Gemmell et al. 2020). Some thermosensitive TRP genes under positive selection in *G. japonicus* were identified, suggesting the important roles of thermoregulation in adaptive evolution of the Schlegel’s Japanese gecko.

In this study, we explored the complex and fascinating evolutionary trajectory of *G. japonicus* based on the chromosome, genome and gene levels, providing novel insights into species-specific and chromosome-specific evolution, adaptive evolution of important traits (e.g. bitter taste perception, detox and biosynthetic cytochrome P450, epidermis formation and thermoregulation) in *G. japonicus*.

## Results

### Sequencing, assembly and annotation of high quality *Gekko japonicus* genome

We combined Illumina Hiseq, Pacbio SMRT sequencing and Hi-C to assemble the chromosome-level genome of *Gekko japonicus*. The results obtained from Illumina Hiseq were first used in a k-mer analysis to estimate the genome size, which resulted in 2.50 Gb (Supplemental Fig. S1). For the reads generated by Pacbio SMRT, the initial genome assembly was first obtained using MECAT2 (Xiao et al. 2017), followed by polishing the genome assembly results with subreads using GCpp version 1.9.0 (https://github.com/PacificBiosciences/gcpp) and error correction based on Illumina data using Pilon version 1.22 (Walker et al. 2014). Finally, Hi-C data was mounted using Juicer version 1.6.2 (Durand et al. 2016) to obtain the final genome assembly of 19 pseudochromosomes (Supplemental Fig. S2; Supplemental Table S1), which is concordant with the previous results (Shibaike et al. 2009). The scaffold N50 is 172.24 Mb and the contig N50 is 2043.27 kb, both greatly improved compared to the previous version (Supplemental Table S2). The size of genome assembly in this study is about 2.54 Gb, which is close to the results of k-mer analysis. We performed BUSCO (Bench marking Universal Single-Copy Orthologs) analysis and the results showed that the genome assembly of *G. japonicus* contains 93.8% complete sequences (Supplemental Table S3), which reveals that the final assembly is highly complete. Our assembly has a lower number of scaffolds and contigs compared to previously published results, but a higher N50 and a larger genome size, which suggests that the assembly results in the present study are more continuous and reliable (Table 1).

**Table 1.** Comparison of genome assembly in this study with previous version

| Assembly | This study | Gekko_japonicus_V1.1 |
| --- | --- | --- |
| Assembly approach | WGS and Hi-C | WGS and BAC |
| Sequencing platform | PacBio, Illumina | Illumina |
| Scaffold N50 (bp) | 172236841 | 707733 |
| Contig N50 (bp) | 2043270 | 29574 |
| Total genes | 20503 | 19548 |
| Total sequence length (bp) | 2535908678 | 2490274461 |
| Total repeat size (%) | 69.4 | 48.9 |
| GC (%) | 45.9 | 45.5 |
| Scaffold Number | 146 | 191500 |
| Contig Number | 2157 | 335470 |

We annotated 69.4% and 65.87% of the genome for repetitive elements and transposable elements, respectively, which is much higher than the previous results, most likely due to the fact that we used more software and methods, including TRF version 4.09 (Benson 1999), RepeatModeler version 1.0.11 (Smit and Hubley 2008), RepeatMasker and RepeatProteinMask version 4.0.9 (Price et al. 2005; Tarailo-Graovac and Chen 2009), to make the annotation more complete (Table 1; Supplemental Tables S4, S5).

A total of 20,503 protein-coding genes (Supplemental Table S6) and 55,481 non-coding RNAs (Supplemental Table S7) were annotated. To demonstrate the improvement of gene structure annotation results in this study relative to the previously published data, 1:1 orthologous gene between two versions were searched using OrthoFinder version 2.5.1 (Emms and Kelly 2015) and although there did not seem to be much difference in terms of length distribution plots (Supplemental Figs. S3, S4), gene length (Wilcoxon signed rank test, one tailed, p < 2.2e-16) and protein length (Wilcoxon signed rank test, one tailed, p = 0.0162) in the present study are both significantly higher than the previous annotation. The comparison of genes, transcripts and exons between the two versions also shows better results for the novel annotation (Table 1; Supplemental Table S8), it suggested that we annotated more genes, transcripts, exons and average number of transcripts per gene, longer average exon length as well as average exon length per gene compared to previous version, while the average transcript length (48,109.86 compared to 48,163.10) and average number of exons per gene (12.66 compared to 12.99) were slightly lower. A search of several different gene annotation databases showed that 96.49% of annotated genes have functional annotations (Supplemental Table S9). All these findings indicate that the genome assembly and annotation in the present study are more complete and greatly improved compared to previous version. The main features of the genome are presented in the Circos plot (Fig. 1). In general, the repetitive elements and genes in all chromosomes show an inverse relationship.

**Figure 1.**
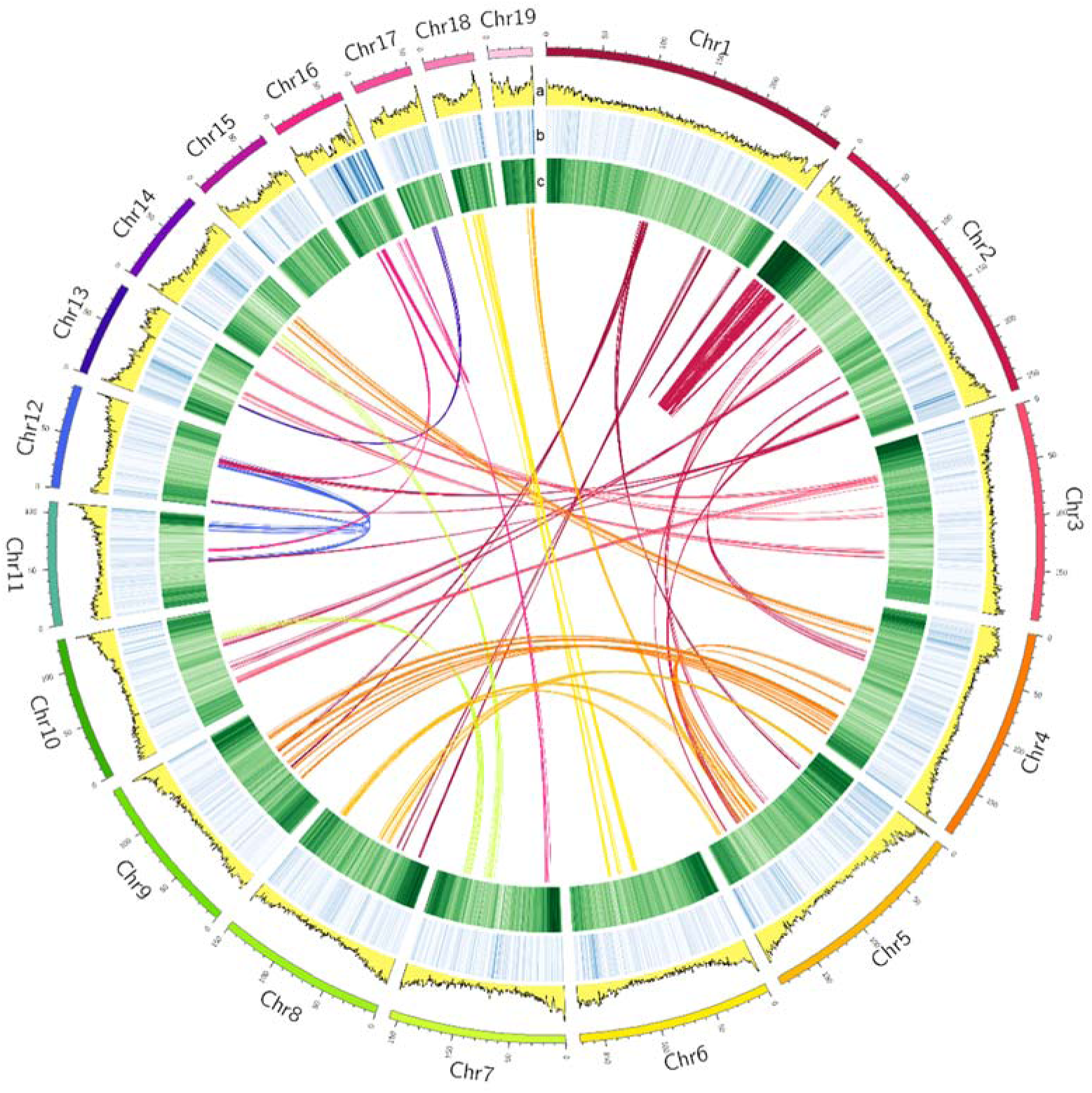
Circos plot of the *Gekko japonicus* genome. The outermost circle indicates the 19 chromosomes information with a tick unit of 10 Mb. Linkages between or within chromosomes show synteny of paralogous genes. The circles labeled with a, b, and c correspond to GC content, gene density, and repeat coverage, respectively, with darker colors representing higher corresponding features.

### Chromosome evolution of *Gekko japonicus*

In order to reveal the specific evolution of chromosomal regions of *G. japonicus*, we selected two high-quality Sauropsidan genomes, *Anolis carolinensis* and *Gallus gallus*, by referring to the identification of corresponding HSBs and EBRs in three Carnivores (Fan et al. 2019). They were aligned with the chromosome sequences of *G. japonicus* and corresponding HSBs were identified. A total of 39 HSBs between *G. japonicus* and *A. carolinensis* and 136 HSBs betwwen *G. japonicus* and *G. gallus* were identified (Fig. 2; Supplemental Tables S10, S11). Synteny analysis showed many chromosomal rearrangement events between *G. japonicus* and the two species, mainly translocations and, to a lesser extent, inversions. In *A. carolinensis*, except for the too short microchromosomes, chromosomes 1 to 6 correspond to at least two chromosomes of *G. japonicus*, and most of the chromosomes of *G. japonicus* have a large segment of colinearity with only one chromosome of *A. carolinensis*, suggesting possible chromosome fission and fussion events from the ancestor of *G. japonicus* to *A. carolinensis*. More rearrangement events have occurred between *G. japonicus* and *G. gallus*, and these results indicate the diversity and complexity of sauropsids chromosome evolution.

**Figure 2.**
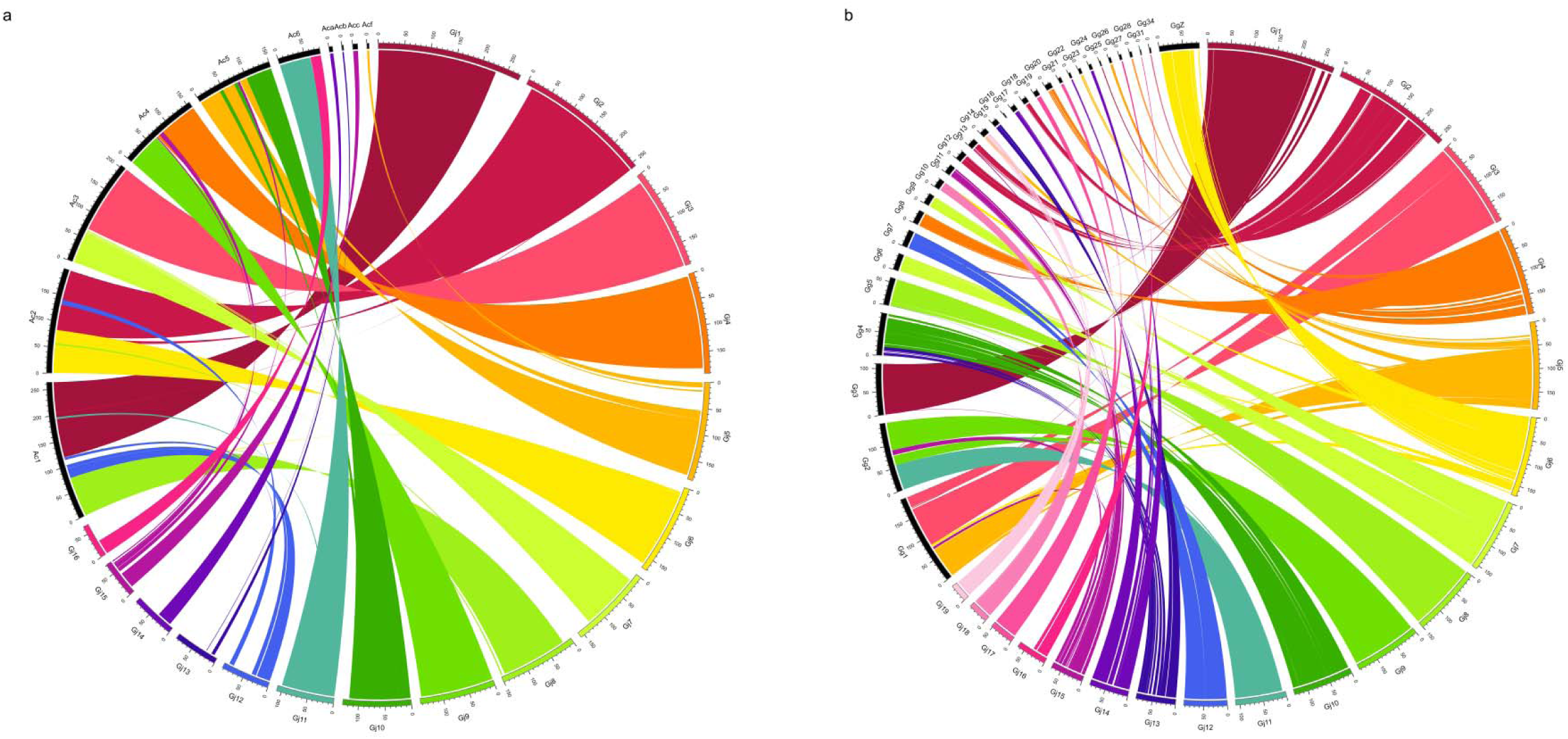
Synteny analysis of *Gekko japonicus* and two other sauropsids. “Gj”, “Ac” and “Gg” are abbreviations of *G. japonicus*, *Anolis carolinensis* and *Gallus gallus*, respectively, and the numbers following them to the right correspond to the respective chromosome numbers. The circle indicates the chromosomes information with a tick unit of 10 Mb. **a,** Circos plot based on the results of synteny analysis between *G. japonicus* and *A. carolinensis*. The chromosomes of *G. japonicus* are marked in non-black and those of *A. carolinensis* in black, and the lines represent the synteny between the corresponding chromosomes. **b,** Circos plot based on the results of synteny analysis between *G. japonicus* and *G. gallus*. The chromosomes of *G. japonicus* are marked in non-black and those of *G. gallus* in black, and the lines represent the synteny between the corresponding chromosomes.

A total of 13 EBRs were identified in *G. japonicus* by HSBs between *G. japonicus* and *A. carolinensis*, with a total length of 106.15 Mb, accounting for 4.24% of the total chromosome size (Supplemental Table S12), while 63 EBRs were identified in *G. japonicus* by HSBs between *G. japonicus* and *G. gallus*, with a total length of 93.83 Mb, accounting for 3.74% of the total chromosome size (Supplemental Table S13). By combining the information of EBRs from two alignment results, a total of 64 merged EBRs were obtained, with a total length of 188.85 Mb, accounting for 7.54% of the total chromosome size (Supplemental Table S14). The length of EBRs in *G. japonicus* ranged from a minimum of 16.97 kb to a maximum of 37.64 Mb, with a mean length of 2.95 Mb and a standard deviation of 6.62 Mb, which indicates that the length of EBRs regions are very variable.

To study the variability of genomic features between EBRs and genomes, we compared gene density, GC content and repeated sequence content across chromosomes between EBRs and genomes (Supplemental Table S15), showing that EBRs have significantly higher gene density (Wilcoxon rank sum test, p = 0.026) and slightly lower repeat content compared to genomes (Wilcoxon rank sum test, p = 0.075 < 0.1), while GC content is not significantly higher in EBRs (Wilcoxon rank sum test, p = 0.76). The results of linear regression (Supplemental Fig. S5) showed an opposite linear relationship between gene density and repeated sequence content in EBRs (adjusted R-squared = 0.237, p = 8.62e-06), which is concordant with higher gene density and slightly lower repeated sequence content in EBRs. We also found that some repetitive elements are significantly higher in EBRs than in the rest of the genome (Supplemental Table S16). These repetitive elements are mainly transposable elements, including DNA transposons (CMC, hAT, Kolobok and PIF), LINE (CR1, L1, L2 and RTE), LTR (ERV1 and Gypsy) and SINE (MIR, tRNA and 5s-Deu-L2), and satellite sequences are also found to be significantly enriched in EBRs, suggests that they may played an important role in chromosome evolution. Enrichment analysis of genes located in EBRs showed that these genes are mainly enriched to defense response pathway (Supplemental Fig. S6), and these immune-related genes, may play an important role in tail regeneration, damage repair, and defense against attack in *G. japonicus*.

### Phylogenetic analysis and evolution of gene families

To reveal the evolutionary relationships between sauropsids and the more accurate phylogenetic position of *G. japonicus* and to clearly show the impact of the assembly of two different versions of the genome on comparative genomics analyses, we selected a total of 18 sauropsids as representatives, one amphibian species and two mammals as outgroups, a total of 22 genomes including the previous version of the *G. japonicus* genome, constructed a phylogenetic tree and estimated the divergence time of each node and performed gene family analysis on them (Fig. 3). The results show that the clade containing *G. japonicus* diverged from other squamate species 191.9 million years ago (Fig. 3a), which is consistent with the fossil record (Daza et al. 2014) and previous research (Liu et al. 2015a). The two versions of *G. japonicus* showed some variability in divergence times, gene family expansion and contraction, and gene clustering analysis, suggests that assembly and annotation with the previous older version may have caused bias in the analysis results, and the importance of high-quality annotation and assembly for comparative genomics analysis. A total of 7408 gene families are conserved in *Anolis carolinensis*, *Alligator sinensis*, *G. japonicus*, *Python bivittatus* and *Pelodiscus sinensis* (Fig. 3b). These gene families may constitute the core proteome of sauropsids. We performed GO and KEGG enrichment analyses for genes in *G. japonicus* that are specific, in expanded gene families, and under positive selection (Supplemental Figs. S7-S11), respectively. The results showed that some of the enriched pathways are associated with sensory and nervous systems such as smell and taste, and that these pathways may be involved in activities such as predation, feeding, and escape from natural enemies, and suggest that their associated genes may have undergone adaptive evolution in *G. japonicus*. Some other pathways are immune-related, such as those related to T cells, Nod-like, and TNF, which play important roles in immune regulation, inflammatory responses (Chen and Goeddel 2002; Chen et al. 2009; Mathew et al. 2009; Platnich and Muruve 2019) and may be important for damage repair, tail regeneration, and defense against pathogens in *G. japonicus*.

**Figure 3.**
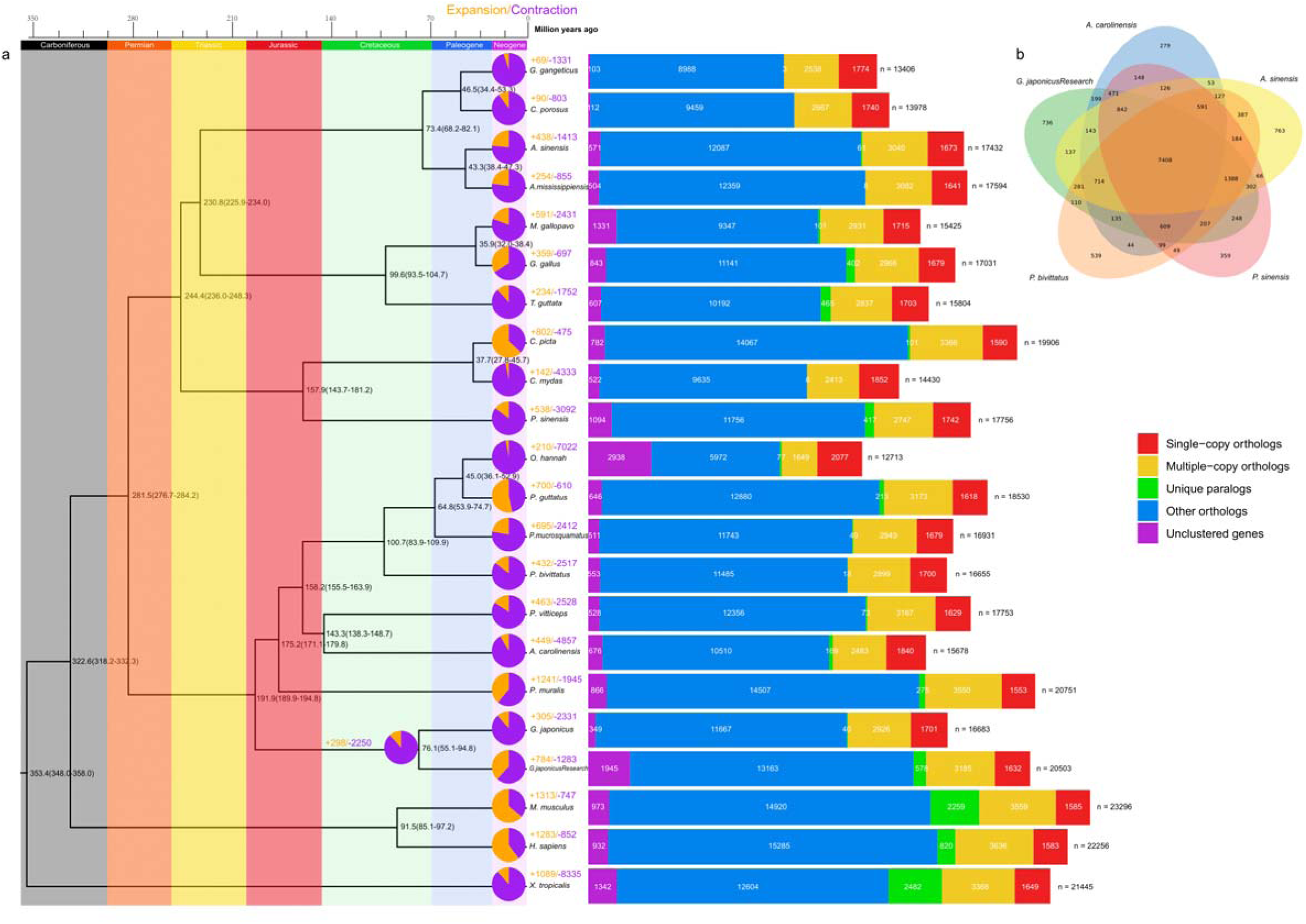
Phylogenetic and gene family analysis of 18 sauropsids and outgroups. “*G. japonicus*” and “*G. japonicusResearch*” correspond to the previous version and the genome assembly of this study, respectively. **a** Phylogenetic tree for 18 sauropsids and outgroups. Bootstrap value for all branches is 100. The numbers on each node and in parentheses represent the estimated divergence time (million years ago, MYA) and the corresponding 95% confidence interval, respectively. Pie charts represent the proportion of gene family expansions and contractions, with purple corresponding to expansions and orange to contractions, and colored numbers corresponding to each of them. The genes of each species are divided into five categories, the colors of which correspond to those in the legend on the right, and are shown in the bar chart, while the white numbers inside represent the number of genes in the corresponding category, and “n” represents the total number. **b** Venn diagram of the shared and unique gene families among six sauropsids. Text and numbers represent corresponding species and the number of gene families, respectively.

### Bitter taste perception and its evolution

Bitterness perception can detect potentially toxic compounds and thus avoid poisoning, which is considered to be an important factor for survival (Behrens et al. 2017). The taste receptor type 2 (T2R) family, a subfamily of the taste receptor family with seven transmembrane segments, plays an significant role in bitter taste perception (Bachmanov and Beauchamp 2007). Its numerical size and gene repertoire composition vary considerably among vertebrates, as evidenced by recent detailed gene identification and phylogenetic studies of the squamates and verterbrates, and demonstrate a significant association between diet and the number of T2Rs (Li and Zhang 2014; Zhong et al. 2019). We collected the T2R protein sequences of western clawed frog, human, mouse and chicken from NCBI (Supplemental Table S17) and used them as reference sequences to identify the T2R sequences of other species. In total, 121 T2R genes were identified, and the number of T2Rs in these seven species was similar to recent studies (Li and Zhang 2014; Zhong et al. 2019), but with some differences, which are controllable considering the differences in methods. The number of T2Rs expanded considerably in the insectivorous *A. carolinensis* and *G. japonicus*, and is significantly greater than in other species, and multiple independent duplications of the T2R in several species were clearly observed in the phylogenetic tree, dispersed among different clades (Fig. 4a). To explore the evolutionary pressures on T2R among these sauropsids, PAML version 4.7 (Yang 2007) and HyPhy version 2.5.2 (Kosakovsky Pond et al. 2020) were used to perform selection pressure analysis. Using the aBSREL (Smith et al. 2015) model to detect branches under positive selection in the whole phylogenetic tree (Fig. 4a), we found that the branches under positive selection were mainly present in *A. carolinens* (13 branches) and *G. japonicus* (4 branches), with only one branch associated with *S. punctatus*, which is consistent with the duplication, expansion and rapid evolution of T2R in the insectivorous *A. carolinens* and *G. japonicus*. Four pairs of site models in PAML and two site models in HyPhy were used to detect positively selected sites (Supplemental Tables S18-S20). The likelihood ratio tests (LRTs) for all four pairs of PAML models were significant (chiq-square test, p < 0.05), considering that the positively selected sites identified by M3 were not recommended by the authors of the software and therefore were not included in the results. The average ω value obtained from M0 is 0.54, which is a relatively high value, indicates that T2R is under relatively low purifying selection on average among these species. There are 9 overlapping sites identified between M2 and M8, and 6 between FEL (Kosakovsky Pond and Frost 2005) and FUBAR (Murrell et al. 2013), but there are few overlaps in the positive selection sites calculated by different software, with only 2 identical sites existing between FEL and M2 or M8 (Fig. 4b). Careful review of the sequence alignment results suggests that this is mainly due to the fact that, in addition to the variability in the calculation methods of the models, PAML removes all sites containing gaps, while HyPhy retains them. Combining the results obtained from all models, a total of 38 positively selected sites were identified, with the most in the extracellular region (ER), 17 (44.74%), followed by the transmembrane region (TM), 15 (39.47%), and the least in the intracellular region (IR), 6 (15.79%), with no significant heterogeneity in this distribution pattern (chi-squared test, chi-squared = 5.42, p = 0.067). Specifically, when T2R was divided into 15 intervals according to the order of ER, TM, and IR from N-terminal to C-terminal, the ER3 region have the most positively selected sites with 6, and the least was IR1, where no positively selected sites were detected (Fig. 4c). In order to detect whether the evolutionary pressure is significantly different in various regions, the pairwise ω values of the ER, TM, and IR region alignments were calculated using YN00 model, respectively, and the average ω was 1.39, 0.89, and 0.45, respectively. The pairwise ω values of the ER were significantly higher than those of the TM (two-sample independent t-test, t = 14.80, p < 2.2e-16) and IR (two-sample independent t-test, t = 16.93, p < 2.2e-16), while the pairwise ω values of the TM region were significantly higher than those of the IR region (two-sample independent t-test, t = 12.39, p < 2.2e-16). These results suggest that the evolutionary pressures on the seven sauropsids T2Rs differ by region, with the ER region being more favoured by positive selection and the TM and IR regions being relatively more constrained by purifying selection, but the TM being less constrained.

**Figure 4.**
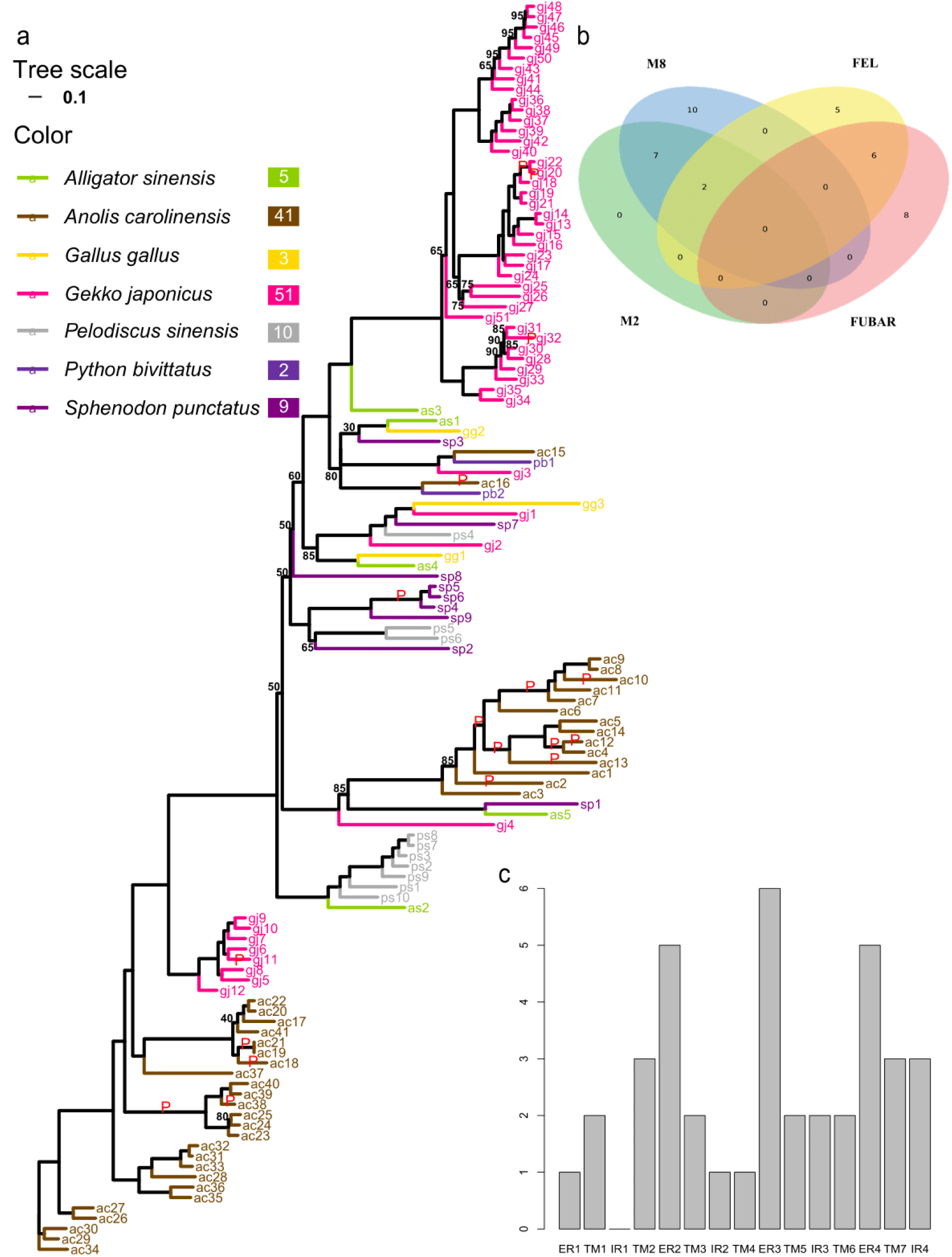
Phylogenetic and selection pressure analysis of the T2R gene in sauropsids. **a** Phylogenetic tree of T2R genes in seven sauropsids. Scale bar for branch length of the tree and the corresponding color of the species are labeled in the legend, and the number to the right of the species name represents the number of T2Rs. Bootstrap values not equal to 100 are labeled by numbers at each node. Branches marked with “P” represent branches that are tested as under positive selection using the aBSREL model. **b** Venn diagram of common and unique positively selected sites in the T2R gene of seven sauropsids identified by four methods. Text and numbers represent the corresponding method and the number of positively selected sites, respectively. **c** Bar graph of the number of positively selected sites in different regions of the T2R gene in seven sauropsids. X-axis corresponds to different regions of the T2R gene, and Y-axis represents the number of positively selected sites in the corresponding regions.

We next focused on the location, phylogeny and evolution of T2R in *G. japonicus*. Among the 51 T2Rs of *G. japonicus* identified, only two (gj1 and gj2) are located on chromosome 1, one (gj3) on chromosome 3 and 48 (gj4-gj51) on chromosome 6, and most of them tandemly clustered (Fig. 5a). It can be found that gj5-gj12 is tandemly clustered on the reverse strand and gj13-gj50 on the forward strand, these two gene clusters correspond to two clades on the phylogenetic tree, while gj51 is close to gj50, but on the reverse strand, and is on the same clade with gj13-gj50 on the phylogenetic tree. All of these results suggest that the expansion of T2R genes is attributed to tandem gene duplication. To investigate the intraspecific evolution of T2R in *G. japonicus* and to compare it with the overall evolutionary pattern of sauropsids, we extracted a subtree of *G. japonicus* from the T2Rs phylogenetic tree of seven species (Fig. 5b) and performed the same evolutionary analyses. A total of six branches under positive selection were detected by the aBSREL model (Fig. 5b), overlapping with the four branches detected with the seven species dataset. These positively selected T2R genes may differ from other T2R genes in bitter taste tolerance, perception and gene expression. The LRT tests for all four pairs of site models performed in PAML were significant (Supplemental Table S21), with the mean ω value calculated by M0 is 0.70, which is higher than the mean ω value of the seven sauropsids, suggests that T2R evolution within *G. japonicus* was more unconstrained by purifying selection. The positively selected sites identified by the FEL and FUBAR models overlapped with four identified by M8 and M2 (Fig. 5c), located at codons 73, 92, 183 and 263 in the multiple sequence alignment, indicating the reliability of these sites. A total of 58 positively selected sites were identified, and their counts in ER, TM, and IR are 23, 24, and 11, with percentages of 41.38, 39.66, and 18.97, respectively, which did not show significant heterogeneity in the distribution (chi-squared test, chi-squared = 5.42, p = 0.067). The distribution of sites divided into 15 regions (Fig. 5d) is consistent with the distribution of the seven species (G-test, G = 33.24, p = 0.77), and a total of nine positive selection sites shared by both were located at codons 13, 66, 87, 88, 90, 154, 162, 240 and 280 in the multiple sequence alignment. The mean ω values of 2.13, 0.68, and 0.44 were obtained from the yn00 analysis for the T2R alignments of the ER, TM, and IR regions, respectively. Similarly, we also found significantly higher mean ω values in the ER region than in the TM region (two-sample independent t-test, t = 4.50, p = 7.3e-06) and the IR region (two-sample independent t-test, t = 8.47, p < 2.2e-16), while the mean ω values in the TM region are significantly higher than in the IR region (two-sample independent t-test, t = 43.80, p < 2.2e-16). The above results clearly indicate that the pattern of evolutionary pressure exerted on *G. japonicus* and that exerted among the seven sauropsids is consistent overall, with both ER and TM being significantly more unconstrained by purifying selection than the IR region, especially the ER region, which has very large levels of inter- and intra-species variation, indicating its importance in adaptive evolution.

**Figure 5.**
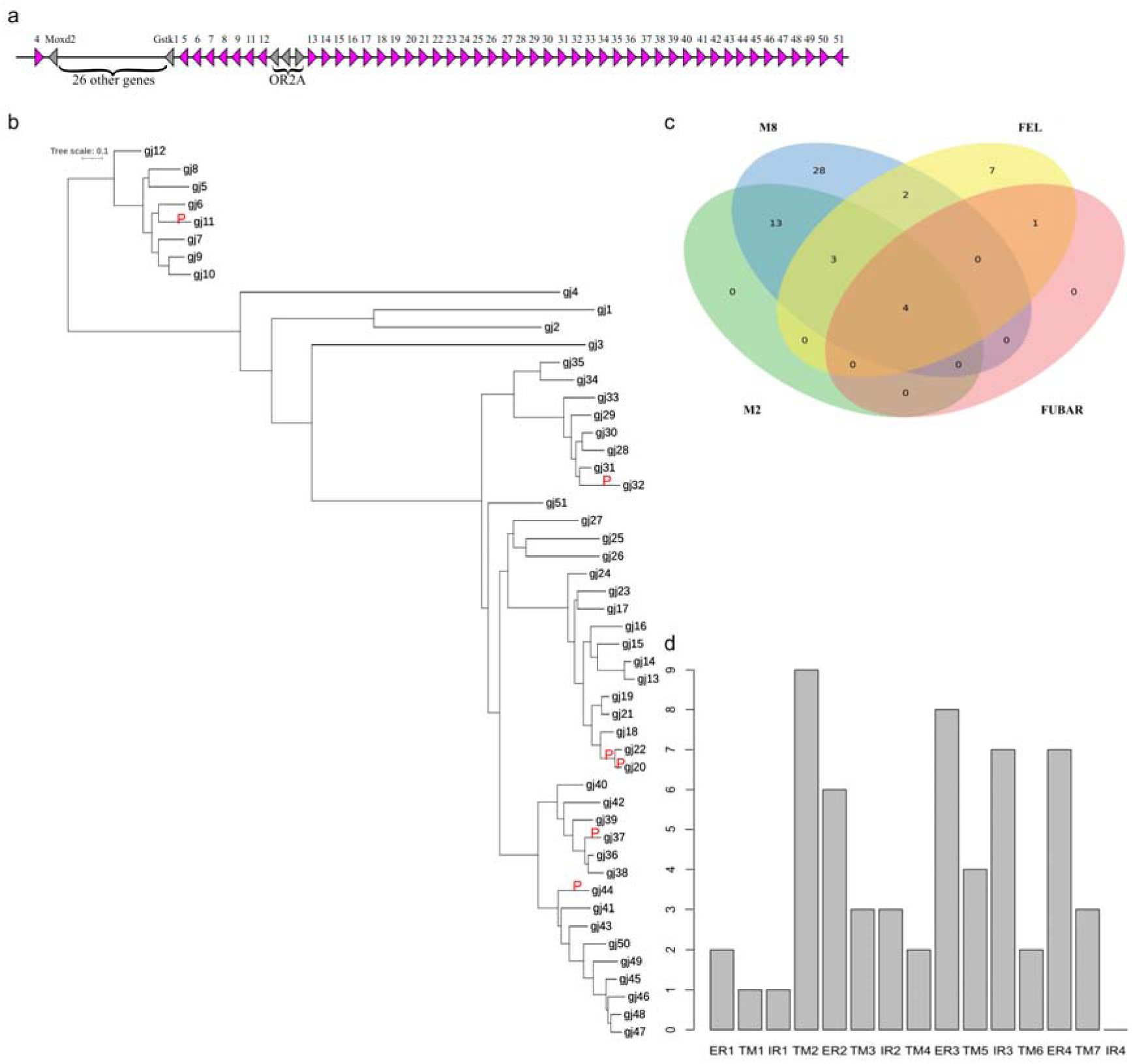
Location and selection pressure analysis of the T2R gene in *G. japonicus*. **a** All 51 T2R genes clustered on chromosome 6 of *G. japonicus*. Red triangles represent T2R genes, gray triangles represent other genes, the obtuse angle direction to the right means in the forward strand, to the left means in the reverse strand. **b** Subtree of the T2R gene in *G. japonicus*. Scale bar of branch length is marked in the upper right corner, and branches marked with “P” represent branches detected as under positive selection using the aBSREL model. **c** Venn diagram of common and unique positively selected sites in *G. japonicus* T2R genes identified by four methods. Text and numbers represent the corresponding method and the number of positively selected sites, respectively. **d** Bar chart of the number of positively selected sites in different regions of the *G. japonicus* T2R genes. X-axis corresponds to different regions of the T2R gene, and Y-axis represents the number of positively selected sites in the corresponding regions.

### Birth and death evolution of CYPs in sauropsids

CYPs are monooxygenases responsible for oxidative and metabolic reactions whose substrates are a range of xenobiotic and endogenous compounds, including various toxic substances. In the three phases of detoxification, CYPs are involved in phase I metabolism, responsible for the oxidation of toxic substances, the oxidized products are then passed to phase II enzymes (e.g., Glutathione S-transferases) for binding and catalysis, and finally removed from the cell via ATP-binding cassette transporters in phase III (Tan and Low 2018). Among sauropsids, many species are exposed to environments where toxic substances are present, such as various insectivorous lizards, geckos and birds. In addition to detecting bitter toxins in insects through the above-mentioned T2Rs, the ingested toxins may be cleared by a series of detoxification genes such as CYPs.

In order to study the birth and death process of CYPs and the evolution of detoxification system in sauropsids, we identified CYPs of 13 representative sauropsids using human functional CYPs as reference. The sauropsids CYPs clustered well into 10 clans, and 18 gene families, but among the subfamilies, phylogenetic relationships and clades divided by MIPhy (Curran et al. 2018) for the phylogenetic tree indicated many sauropsids CYPs are not orthologs to human CYPs and have different subfamily numbers from human CYPs or are difficult to assign to existing subfamilies of human CYPs (Fig. 6; Supplemental Fig. S12). CYP1B has two subclades, one of which has genes that do not cluster with human CYP1B1 and is therefore named CYP1B2. CYP2F has three subclades, two of which have genes that do not cluster with human CYP2F1 and are therefore named CYP2F2 and CYP2F3. CYP2W has two subclades, one of which has genes that do not cluster with human CYP2W1 and is therefore named CYP2W2. CYP4B is similarly divided into two subclades. CYP3A has three sub-clades, CYP3A9 and CYP3A24 named by the results of the blastp search, and the separately clustered human CYP3A. A unknown clade named CYP2H1, which does not cluster with any CYP2 gene family. Consistent with expectations, the number of D-type CYP genes is significantly more variable compared to B-type CYP genes (Supplemental Fig. S13; Supplemental Table S24), and MIPhy scores are significantly higher in D-type than in B-type CYP genes (two-sample independent t-test, t = 3.1125, p-value = 0.00469), the results indicate that the clades of D-type CYP genes has higher instability. D-type CYP genes have frequent duplication and loss events, suggests that they undergo more birth and death processes compared to B-type CYP genes (Supplemental Table S25).

**Figure 6.**
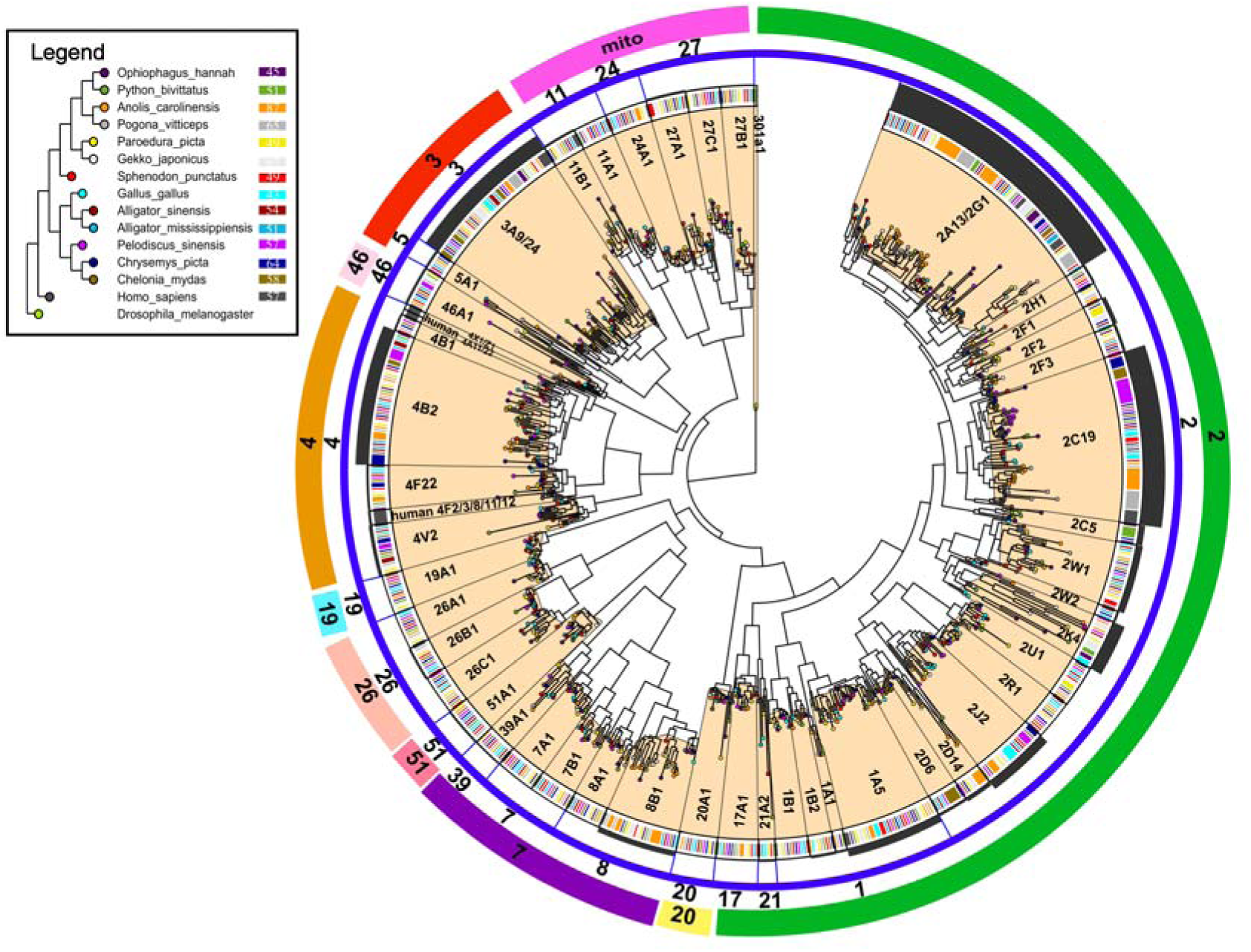
MiPhy analysis and phylogeny for sauropsids P450. The numbers correspond to clan, gene family and specific gene name from the outside to the inside of the circle. The width of each black circle arc corresponds to the size of the MIphy score, and the color of the endpoints in the tree corresponds to the species as shown in the legend in the upper left corner, and the numbers in the legend indicate the size of the number of genes in the corresponding species.

We calculated ω values for each clade divided by MIPhy using the M0 model in PAML (Supplemental Table S26) and found a correlation between ω values and MIPhy scores (Spearman’s correlation, ρ = 0.37, p = 0.012), however, no significant differences were detected in ω values between D-type and B-type CYP genes (average ω 0.18 vs. 0.15, two-sample independent t-test, t = 1.2011, p-value = 0.2363), although the mean ω value is slightly higher in D-type CYP genes. Positively selected sites were found in 13 of the 46 clades using M2a and M8 model in PAML (Supplemental Table S26), including 6 from 24 D-type related clades and 7 from 22 B-type related clades. 15 branches were found to be under positive selection, 12 of which were from clades of D-type CYP genes (Supplemental Table S27). CYP2D14, CYP2F3, CYP2U1, CYP4F22 and CYP26C1 were found under positive selection in foreground branches of *G. japonicus*, but only CYP2D14 was significant after multiple hypothesis testing correction.

To investigate whether dietary differences are related to changes in the number of CYPs, we performed phylogenetic independent contrast (PIC) analysis of CYP1-CYP4 clades, as most of these clades are unstable, have frequent birth and death processes and vary somewhat among different species. However, we only found a correlation between CYP1 and diet (Supplemental Fig. S14). These results demonstrate the complexity of the interaction between selection pressure and birth-death processes.

### Phylogeny, structural composition and synteny of CBPs, an important gene family related to epidermis formation

Epidermal differentiation is one of the most significant phenotypic changes for amniotes during the evolution from aquatic to terrestrial ecosystem. Epidermal appendages, e.g. feathers, claws, scales, etc., are composed of keratins and other relative proteins, especially the corneous beta proteins (CBPs). CBPs are special proteins that expressed in Sauropsida only and coded by the CBP genes family. The CBP genes family has been identified in different animals such as birds (Greenwold and Sawyer 2010), lizards (Dalla Valle et al. 2010; Hara et al. 2018; Holthaus et al. 2020), snakes (Dalla Valle et al. 2007; Alibardi 2014), crocodilians (Alibardi and Thompson 2002; Alibardi and Toni 2007; Holthaus et al. 2018), turtles (Holthaus et al. 2016; Alibardi 2020), etc. Previous researches have identified various numbers of the CBP genes family in several different animals. For example, 111 complete CBP genes sequences (coding regions) and a total of 108 β-keratin genes were identified in the *Gallus gallus* and *Taeniopygia guttata* genome respectively (Greenwold and Sawyer 2010), 89, 37, and 74 beta-keratin genes in the genomes of three turtle species (*Chrysemys picta*, *Chelonia mydas*, and *Pelodiscus sinensi*) (Li et al. 2013), 40 in *Anolis carolinensis* (Dalla Valle et al. 2010), 71 in *Gekko japonicus* (Liu et al. 2015b), 35 and 36 in *Python bivittatus* and *Ophiophagus hannah* respectively (Holthaus et al. 2017), 22 in *Alligator mississippiensis* and *Crocodylus porosus* (Holthaus et al. 2018), and 36 in *Sphenodon punctatus* (Holthaus et al. 2020). Various numbers of the CBP genes in different species may be caused by duplication or loss of these genes. However, the specific gene locations and their functions have not been studied, therefore we next conducted a comprehensive study of CBPs in sauropsids.

The identified CBP genes sequences in *Gekko gecko* (Hallahan et al. 2009) and *Anois carolinensis* (Dalla Valle et al. 2010) were used as reference to search porteins of selected reptiles and others (see Methods for more details). 66 amino acid sequences of CBP genes family members of *G. japonicus* were obtained (Supplemental Table S29). These sequences were tentatively named CBP1 ~ CBP66 according to their positions on chromosomes. The CBP genes sequences of *G. japonicus* were principally located in the EDC region of chromosome 1. In this study, the CBP genes of 12 reptiles (the CBP genes family have been studied) and six reptiles, western clawed frog and mouse (the CBP genes family have not been broadly studied) that with good genome assembly and annotation data were identified (Table 2). Among them, the number of CBP genes families of some species identified in this study is generally less than that have been published. Except, the number of CBP genes family of *Paroedura picta* is the same as that of existing studies, which is 120, and the number is the largest among the species studied so far. However, in *Eublepharis macularius* and *Shinisaurus crocodilurus*, the present study identified more numbers than that of have been reported. By comparison, the number of CBPs in Crocodilian is the least among reptiles, with an average of 9. Among squamates, there are 10 CBPs sequences on average in snakes and 38 in lizards. The average number of CBPs in Chelonia is 18. No CBP genes family has been identified in *Mus musculus* and *Xenopus tropicalis*. The difference in the number of CBPs members may be related to the characteristics of their different epidermis.

**Table 2.**
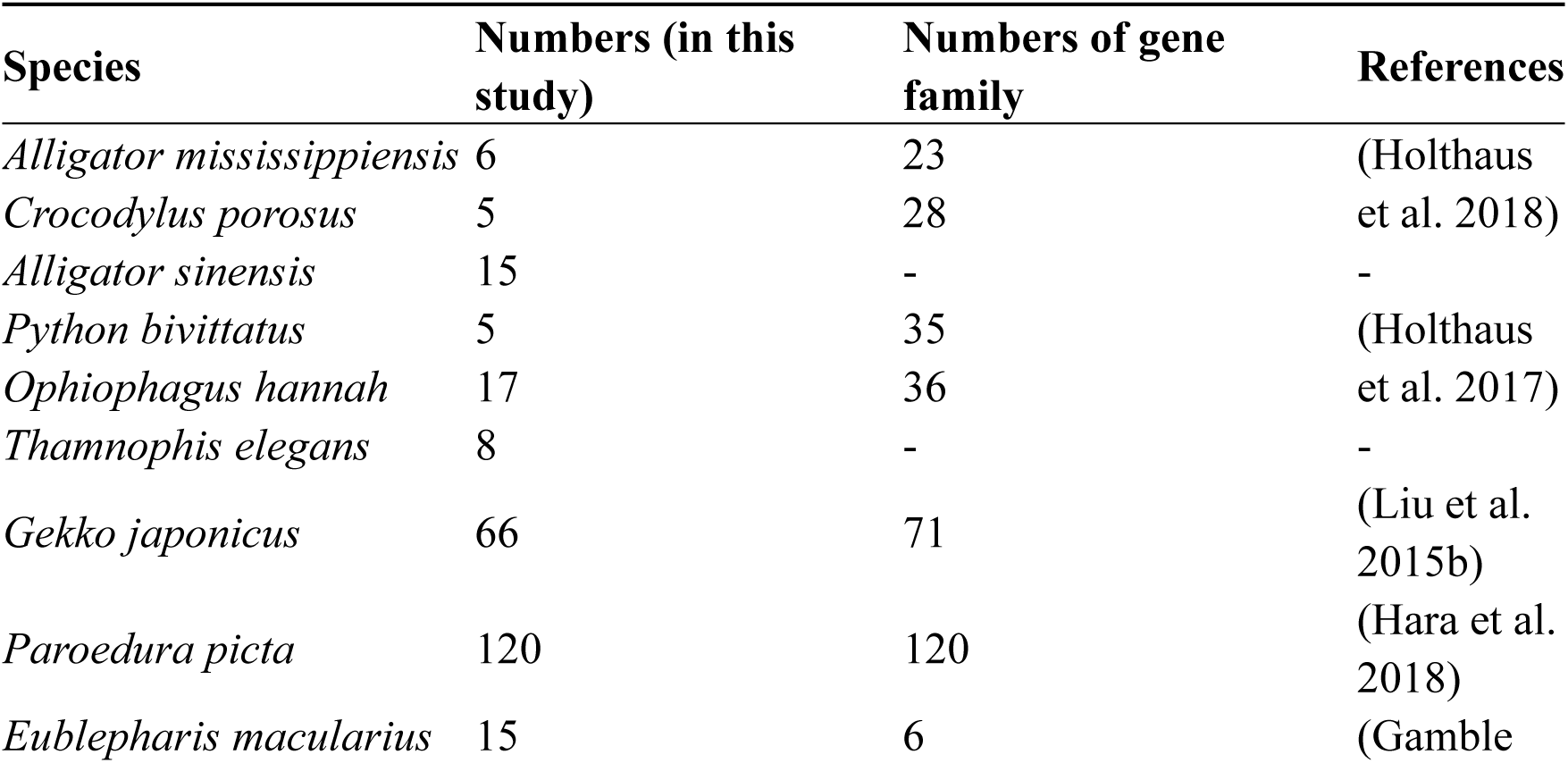
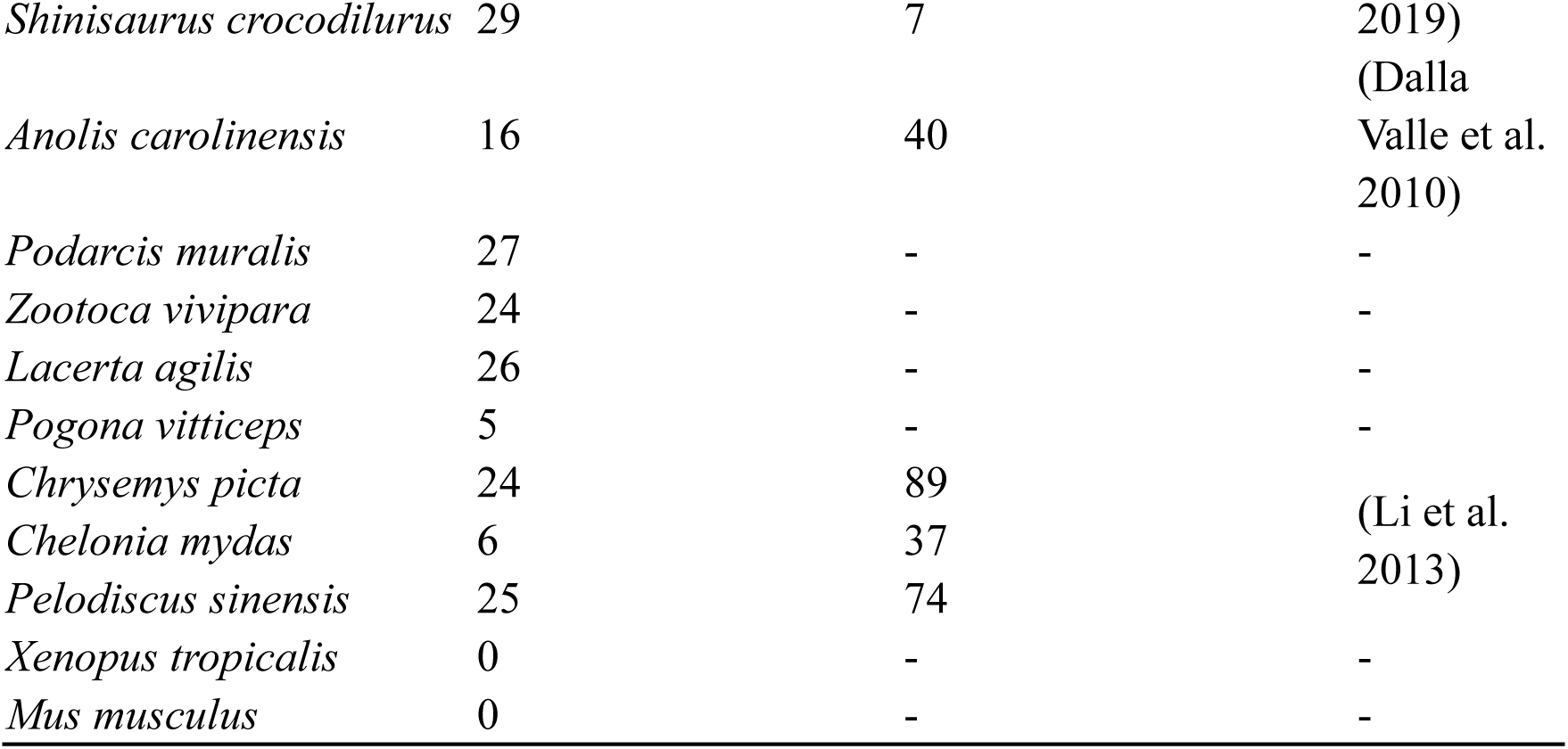
Comparison of the number of CBP gene families in 18 reptiles and mouse

Among the 66 amino acid sequences of CBP genes family in *G. japonicus*, the shortest length was 85 aa, the longest was 478 aa, and the average length was 151.73 aa. The minimum molecular weight was only 8.92 kDa, and the maximum was 41.88 kDa, with an average of 15.23 kDa. It can be seen that these CBPs proteins are all small proteins with short sequence length and small molecular weight. The theoretical isoelectric point values range from 4.14 to 10.26. Although the data vary widely, most of them range from 7.0 to 9.0, and there are more neutral and alkaline sequences. Proline (Pro), cysteine (Cys), glycine (Gly) and serine (Ser) were the most abundant amino acid sequences in 66 CBPs sequences of *Gekko japonicus* (Supplemental Fig. S15). The highest contents of Pro, Cys, Gly and Ser are 22%, 23%, 44% and 22% respectively. In general, these proteins contain small amounts of charged amino acids such as glutamic acid (Glu), aspartic acid (Asp), arginine (Arg), and lysine (Lys).

Align of 66 amino acid sequences of CBPs in *G. japonicus* (Supplemental Fig. S16) revealed that each sequence contained a very conserved motif known as “core box”. The motifs of 66 amino acid sequences of CBPs in *G. japonicus* were predicted (Supplemental Fig. S17). Each sequence contained a very conservative motif 1, which was consistent with sequence alignment, especially the short sequences “PPP” and “PGP” are present in each sequence. Only one sequence (rna-Gja001967.1) contained four motif regions, and three sequences (rna-Gja001974.1, rna-Gja001992.1, rna-Gja002040.1) contained two motif regions. In addition to the conserved motifs, conserved domain analysis of these sequences was performed using the NCBI conserved domain analysis website (Supplemental Fig. S18). 35 of the 66 sequences in the *G. japonicus* contained conserved keratin superfamily domains. However, there has been little analysis of the conserved domain of sauropsids.

Phylogenetic tree of 66 amino acid sequences of CBPs in *G. japonicus* was constructed, and these sequences were divided into three categories. After combining phylogenetic tree results with the motifs, domains and gene structure distribution map mentioned above (Supplemental Fig. S18), it was found that the sequences with similar motif structure basically clustered together. Phylogenetic tree was constructed with CBPs sequences of the selected 20 species (Fig. 7). The CBPs sequences of squamates were basically clustered together. However, these sequences were not completely clustered by species evolutionary relationships, suggests that the CBP genes family may have evolved specifically in each clade.

**Figure 7.**
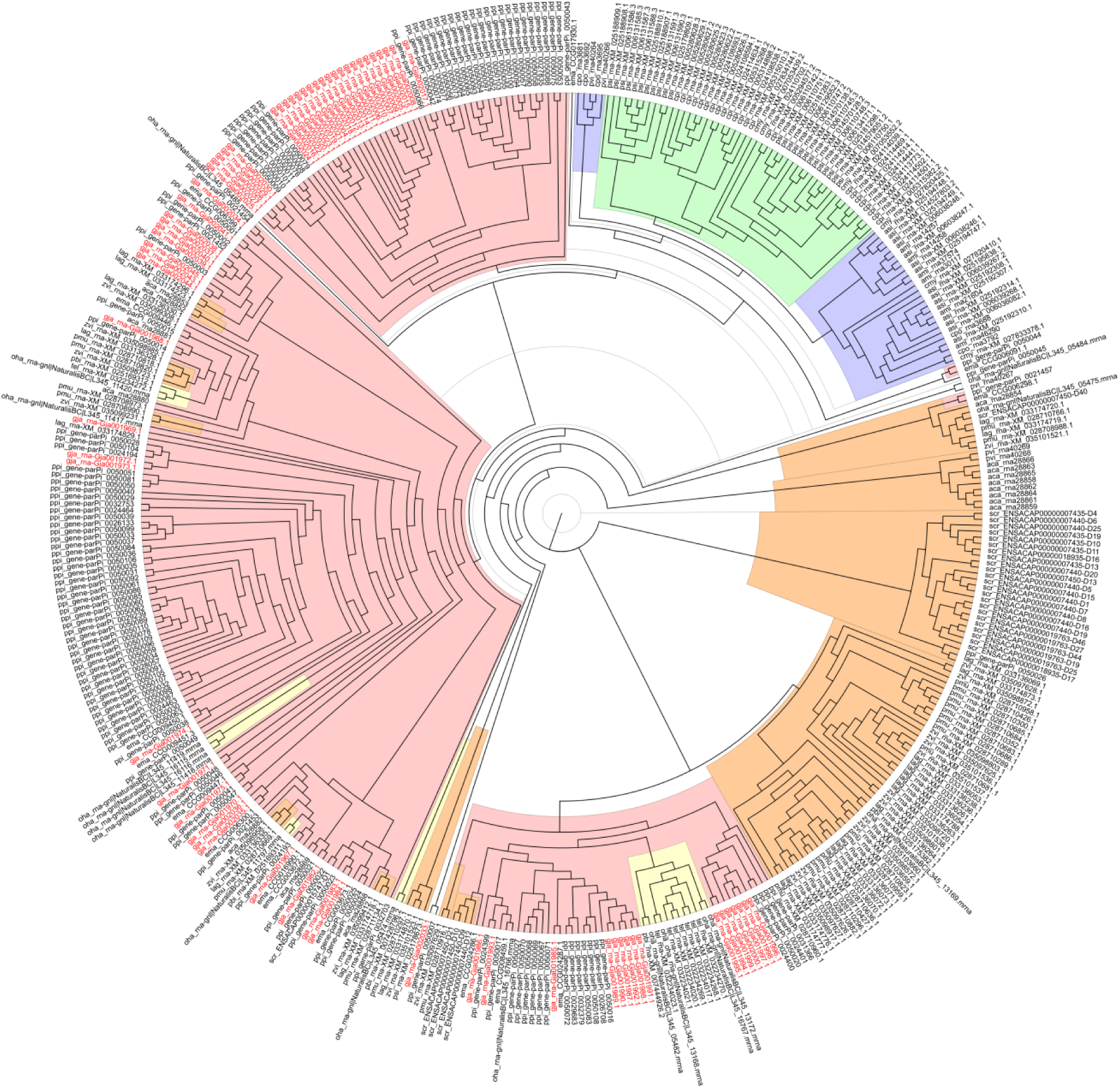
Phylogenetic tree of CBP gene families in 20 species. Red background indicates geckos, orange background indicates lizards, yellow background indicates snakes, green background indicates turtles and crocodiles, and blue background shows crocodiles. CBP sequences of *G. japonicus* are shown in red font.

The arrangement of the CBP genes on the chromosome or scaffold of the selected 20 species (Supplemental Fig. S19) is basically similar, most of which are arranged in series on the same chromosome or scaffold. Besides the arrangement of the CBP genes, there also exists the flanking LOR genes. However, some genomes were not assembled at the chromosomal level, the identified CBP genes in some species were arranged on multiple scaffolds or contigs. Most of the CBP genes family members in the *G. japonicus* have syntenic relationship with that of other reptile species (Supplemental Fig. S20), especially with the *Paroedura picta*, *Podarcis muralis*, *Zootoca vivipara*, *Lacerta agilis*, and *A. carolinensis*. However, they have no syntenic relationship with Crocodilian and Chelonian. This indicates that the CBP genes family of the *Gekko japonicus* is more similar to that of the lizards in evolution.

### Evolution of TRP genes in sauropsids

TRP genes family is involved in a variety of physiological functions, including thermoregulation, reproduction, lysosomal function, inflammatory response, etc., and is widely conserved in vertebrates. It has seven subfamilies, including TRPA1, TRPV, TRPC, TRPM, TRPML (also called MCOLN), TRPP (also called PKD) and TRPN (Clapham et al. 2001; Montell et al. 2002; Yu and Catterall 2004; Clapham et al. 2005). The researches on TRP genes of model organisms have revealed that among the TRP superfamily, TRPA1, TRPM2-5, TRPM8 and TRPV1-4 are thermosensitive (Voets et al. 2004; Wang and Siemens 2015). Some researches on TRP genes in lizards, frogs and crocodiles have revealed the important role of thermosensitive TRPs in the behavior, physiology and thermoregulation of ectotherms, and found that thermosensitive TRPs have different activation temperatures in different species and the activation temperatures of thermosensitive TRPs in reptiles and frogs show an opposite pattern to that of mammals (Seebacher and Murray 2007; Saito et al. 2011; Ohkita et al. 2012; Saito et al. 2012; Kurganov et al. 2014; Saito and Tominaga 2015; Saito et al. 2016; Saito and Tominaga 2017). Even for anole lizards of the same genus, the activation temperature of TRPA1 varies depending on the geographic environment in which they live (Akashi et al. 2018). Here, we selected a total of 16 sauropsids and 3 other species (human, mouse and western clawed frog) as representatives, analyzed their TRP genes repertoire, and attempted to reveal the shift of selection pressure on TRPs in the temperature adaptation of *G. japonicus*.

TRP sequences from human, mouse, chicken and previous versions of *G. japonicus* as a reference were collected to identify TRP sequences of *Sphenodon punctatus*, *Vipera berus*, *Ophiophagus hannah*, *Paroedura picta* and *Gekko japonicus* (this study). The total number of TRP genes identified in *S. punctatus*, *V. berus*, *O. hannah*, *P. picta* and *G. japonicus* were 31, 33, 31, 31 and 31, respectively (Supplemental Table S30). The number of TRP genes family, and the number and composition of each subfamily, are very similar among different sauropsids, suggests that TRPs are conservative in their evolution. Moreover, the constructed phylogenetic tree that referring to recent study rooted with VDAC (Gemmell et al. 2020) (Fig. 8a) indicated the clade size of each subfamily is very similar, which is due to the fact that each subfamily has almost only one copy of its submember among sauropsids, with multiple copies in a few species. In *G. japonicus*, we found no TRPM6, PKD1L1 and TRPN, and only one copy was found for all other TRP subfamily members. PKD1L1 is missing in most sauropsids, with a single copy present only in *C. mydas* and *S. punctatus*. PKD1L3 and TRPN are missing in some sauropsids, and are absent in all researched snakes. Whether these gene losses are specific to the corresponding lineage requires more high-quality genome samples of representative species for comprehensive analyses and confirmation.

**Figure 8.**
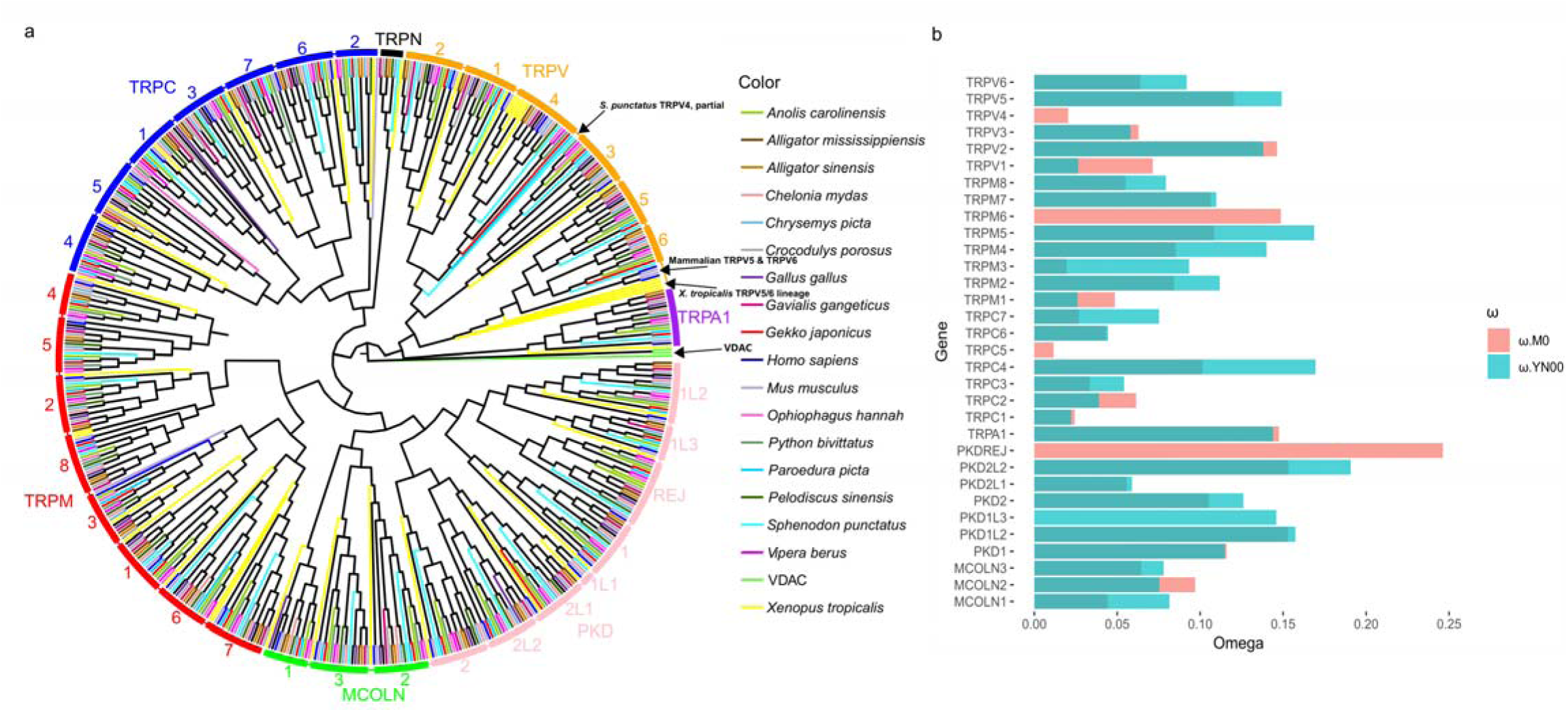
Phylogenetic tree of the TRP gene family and the ω values of each gene. **a** Phylogenetic tree of TRP genes in 16 sauropsids, 2 mammals and 1 amphibian. The outer circle is divided by different TRP subfamilies and their submembers, with the same subfamilies labeled with the same color and the submembers labeled with text. Different colored branches correspond to different species, as shown in the legend on the right. **b** Bar graph of ω values of different TRP genes. X-axis represents the ω value and Y-axis represents the corresponding gene. ω values identified with M0 and YN00 are labeled with different colors, as shown in the legend on the right.

To test the change and shift of selection pressure on *G. japonicus*, we used the branch-site model and clade model in PAML with *G. japonicus* set as a foreground and investigate the difference in TRPs evolution with other sauropsids. A total of six TRP genes detected shifts in selection pressure, which were TRPV5, TRPM3, TRPM5, TRPM8, MCOLN1 and PKD2L2 (Table 3; Supplemental Tables S31-S62). Most of the best models with the lowest AIC values were CmC and CmD, only in PKD2L2 both CmC and CmD were not significant, and the lowest AIC was M2a_rel. Except for MCOLN1, which showed intensified purifying selection, all the other five TRP genes revealed higher ω in the foreground branch, indicating the rise of ω in *G. japonicus* and the presence of purifying selection relaxation, while TRPM3 and PKD2L2 were detected under positive selection in the foreground branch. However, after correction for multiple hypothesis testing, only the LRT for PKD2L2 was significant (Supplemental Table S63), but divergent selection may still be present in the other five TRP genes and should not be ignored. To effectively detect relaxation of selection pressure in *G. japonicus*, we also used branch model, including paired one-ratio model (M0) and two-ratio model (Supplemental Tables S31-S61). The TRP genes with p-values less than 0.05 obtained from the likelihood ratio tests are TRPM3, TRPM8 and MCOLN1 (Supplemental Table S62), and after correction for multiple hypothesis testing, only TRPM3 has a q-value less than 0.05 (Supplemental Table S63). Except for MCOLN1, which revealed intensification of purifying selection, all other genes showed relaxation of purifying selection in *G. japonicus*, and these results are consistent with the results of the branch-site model and the clade model. We also used four pairs of site models to detect positively selected sites for TRP genes, including M3 vs. M0, M2a vs. M1a, and M8 vs. M7/M8a (Supplemental Tables S26-S56), but it was hard to find positively selected sites with PAML in our data, which also indicates that TRP family members are conserved in general. The ω values calculated by one-ratio model (M0) indicate that the TRP genes in general are mainly under relatively strong purifying selection in sauropsids (Fig. 8b), with an mean ω value of 0.085, SD of 0.053. The gene under strongest purifying selection was TRPC5 with an ω value of 0.012 and the weakest was PKDREJ with an ω value of 0.25. Given the low quality or absence of some TRP genes in a large number of species samples, we did not run the M0 model to calculate mean ω values for them. To complement these results, we selected 1:1 orthologous genes between *A. carolinesis* and *G. japonicus* for YN00 analysis (Fig. 8b). The results obtained by YN00 and those obtained by M0 for the shared genes are not different in general (Kolmogorov-Smirnov test, p = 0.53), indicate that the two results are consistent, although the ω values obtained by YN00 is higher than that of M0 (paired sample t-test, t = −2.76, p = 0.0106). The gene with the lowest ω value calculated by YN00 was TRPC1 with 0.022 and the highest was PKD2L2 with 0.19. PKDREJ and PKD2L2, two important genes associated with reproduction, possess higher ω as expected (Vicens et al. 2015). These results are consistent with many previous studies of reproduction-associated proteins, which have shown that reproduction-associated proteins have higher evolutionary rates than other proteins, that many of them are under positive selection, and that they diverge rapidly between taxa (Swanson et al. 2001; Swanson and Vacquier 2002; Clark et al. 2006; Walters and Harrison 2010; Walters and Harrison 2011; Wilburn and Swanson 2016; Twort et al. 2017).

**Table 3.**
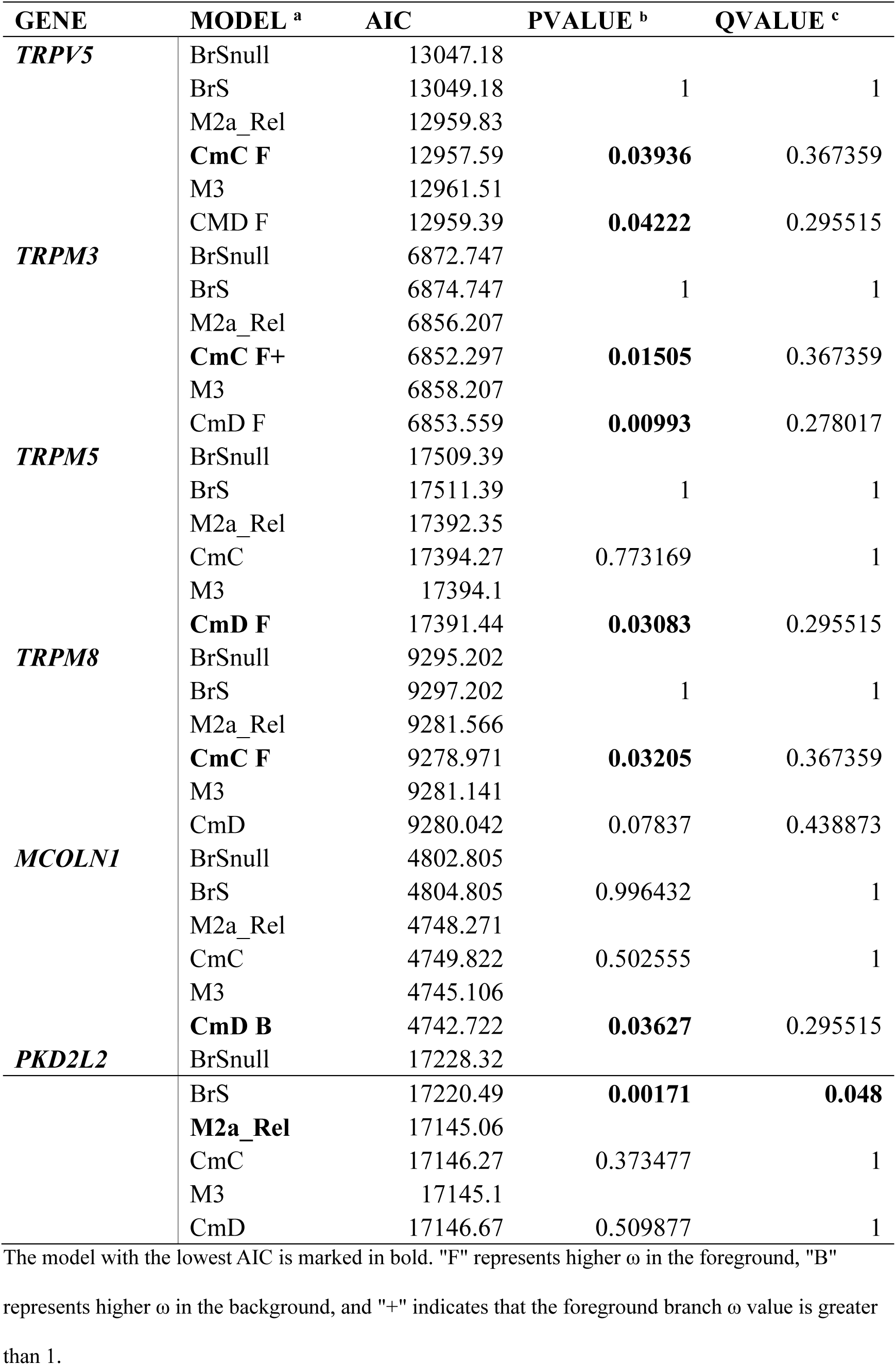
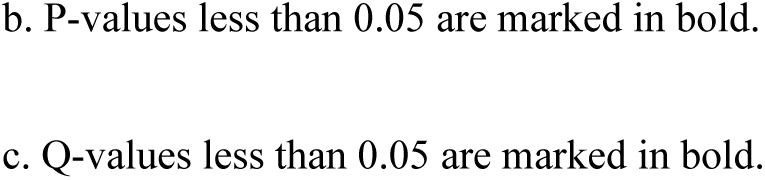
TRP genes that are significant in LRTS of PAML analysis

| GENE | MODEL <sup>a</sup> | AIC | PVALUE <sup>b</sup> | QVALUE <sup>c</sup> |
| --- | --- | --- | --- | --- |
| *TRPV5* | BrSnull | 13047.18 |  |  |
|  | BrS | 13049.18 | 1 | 1 |
|  | M2a_Rel | 12959.83 |  |  |
|  | **CmC F** | 12957.59 | **0.03936** | 0.367359 |
|  | M3 | 12961.51 |  |  |
|  | CMD F | 12959.39 | **0.04222** | 0.295515 |
| *TRPM3* | BrSnull | 6872.747 |  |  |
|  | BrS | 6874.747 | 1 | 1 |
|  | M2a_Rel | 6856.207 |  |  |
|  | **CmC F+** | 6852.297 | **0.01505** | 0.367359 |
|  | M3 | 6858.207 |  |  |
|  | CmD F | 6853.559 | **0.00993** | 0.278017 |
| *TRPM5* | BrSnull | 17509.39 |  |  |
|  | BrS | 17511.39 | 1 | 1 |
|  | M2a_Rel | 17392.35 |  |  |
|  | CmC | 17394.27 | 0.773169 | 1 |
|  | M3 | 17394.1 |  |  |
|  | **CmD F** | 17391.44 | **0.03083** | 0.295515 |
| *TRPM8* | BrSnull | 9295.202 |  |  |
|  | BrS | 9297.202 | 1 | 1 |
|  | M2a_Rel | 9281.566 |  |  |
|  | **CmC F** | 9278.971 | **0.03205** | 0.367359 |
|  | M3 | 9281.141 |  |  |
|  | CmD | 9280.042 | 0.07837 | 0.438873 |
| *MCOLN1* | BrSnull | 4802.805 |  |  |
|  | BrS | 4804.805 | 0.996432 | 1 |
|  | M2a_Rel | 4748.271 |  |  |
|  | CmC | 4749.822 | 0.502555 | 1 |
|  | M3 | 4745.106 |  |  |
|  | **CmD B** | 4742.722 | **0.03627** | 0.295515 |
| *PKD2L2* | BrSnull | 17228.32 |  |  |
|  | BrS | 17220.49 | **0.00171** | **0.048** |
|  | **M2a_Rel** | 17145.06 |  |  |
|  | CmC | 17146.27 | 0.373477 | 1 |
|  | M3 | 17145.1 |  |  |
|  | CmD | 17146.67 | 0.509877 | 1 |

The model with the lowest AIC is marked in bold. "F" represents higher $\omega$ in the foreground, "B"
represents higher $\omega$ in the background, and "+" indicates that the foreground branch $\omega$ value is greater
than 1.
b. P-values less than 0.05 are marked in bold.
c. Q-values less than 0.05 are marked in bold.

## Discussion

Genome assembly at the chromosome level is a valuable resource that allows us to study species-specific evolution from a chromosome perspective. Currently, there are only a few species with chromosomal-level genome assembly among reptiles, and it is difficult to perform more accurate genes identification and evolutionary analyses. Here, we combined Illumina Hiseq sequencing, PacBio SMRT sequencing, and Hi-C technology to assemble a high-quality *G. japonicus* genome with a chromosome number of 19 and a genome size of 2.54 Gb. The genome assembly in this study is more contiguous and higher quality than previous versions, which is reflected in the genome’s assembly higher N50, fewer scaffolds and contigs (Table 1). The higher size of *G. japonicus* genome in the present study may also be due to the greater contiguity. Not only at the assembly level, but also the annotation of the genome has been greatly improved compared to the previous version, with more, complete and detailed annotation results than the previous version. With such high quality genomes, we have been able to conduct comprehensive comparative genomics and evolutionary analyses in combination with other sauropsids.

Chromosome evolution is important for species formation, but comprehensive bioinformatics studies on sauropsids chromosome evolution are scarce. Murphy et al. (Murphy et al. 2005) studied mammalian chromosome evolution as early as 2005 and found that nearly 20 percent of chromosome breakpoint regions were reused during mammalian evolution and that the gene density in EBRs was higher compared to the genome-wide level. Elsik et al. (Elsik et al. 2009) identified cattle-specific EBRs in 2009 and found higher segmental duplications and also found some repeatitive elements enriched in them, mainly LINE-L1 and LINE-RTE. Moreover, Larkin et al. (Larkin et al. 2009) in 2009 identified EBRs from 10 species of amniotes and found that genes related to adaptive function were significantly enriched in EBRs, and also found that retrotransposed genes, zinc finger genes, and CpG islands are significantly higher in EBRs, demonstrating the importance of EBRs in adaptive phenotypes, functional evolution, and species formation. Then, Groenen et al. (Groenen et al. 2012) identified pig-specific EBRs in 2012 and found an enrichment of multiple repetitive elements such as LTR-ERV1, LINE-L2, satellite sequences, and genes located in EBRs enriched to pathways associated with taste perception. Farré et al. (2016) (Farré et al. 2016) identified bird-specific EBRs and found four transposable elements LINE-CR1, LTR-ERVL, LTR-ERVK and LTR-ERV1 enriched in lineage-specific EBRs. Genes located in the lineage-specific EBRs are enriched in several different GO terms. Futhermore, Fan et al. (2019) (Fan et al. 2019) identified HSBs and EBRs specific to each species by pairwise synteny analysis of three carnivores including dog, cat and giant panda, and found enrichment of LINE-L1 elements in the EBRs of the three carnivores, with higher GC content, gene density and repeat elements in the EBRs of giant panda and dog compared to whole genome, while no such differences were found in cat. Some genes located in the EBRs of giant panda, dog and cat are enriched in olfactory perception-related pathways. In this study, we identified the corresponding HSBs and EBRs by pairwise synteny analysis of *G. japonicus* with two other representative sauropsids, *A. carolinensis* and *G. gallus*, which possess relatively high-quality chromosomal-level genome assembly and annotation and differ in chromosome number, following the method of Fan et al (Fan et al. 2019). The specific EBRs in *G. japonicus* have higher gene density and lower repetitive elements compared to the whole genome, which is consistent with the fact that both are in some degree linear and are negatively correlated in EBRs. Howerver, there was no significant difference in GC content between EBRs and the whole genome. DNA transposons (CMC, hAT, Kolobok, and PIF), LINE (CR1, L1, L2, and RTE), LTR (ERV1 and Gypsy), and SINE (MIR, tRNA, and 5s-Deu-L2), and satellite sequences are significantly higher in the EBRs region specific to *G. japonicus* compared to the rest of the genome. The genes located in the EBRs are mainly immune-related genes, significantly enriched in defense response pathway, and these are likely to be important for various immune functions and activities in *G. japonicus*, such as damage repair, tail regeneration, etc. In summary, evolutionary breakpoint regions (EBRs), which resulted from non-random breaks in chromosomes, are essential regions and hotspots for evolution. Specific genes located in EBRs, higher repetitive elements, may have a very strong impact on species evolution and species formation. Of course, our study was limited to single species-specific EBRs and only three species were compared, after which as more and more high quality reptile species, and bird species genomes are assembled to the high quality chromosome level, it becomes possible to sample all taxa of Sauropsida evenly. Performing multi-species, large-sample, comprehensive sauropsids chromosome evolutionary analyses is something that needs to be done in the future.

Phylogenetic analyses and estimated divergence times for representative sauropsids were performed to provide a more precise phylogenetic position. We found some variability in comparative genomics analysis between our genome assembly and the previous one. Gene family clustering, gene family expansion and contraction and positive selection pressure analyses revealed important functional gene sets in *G. japonicas* genome in our assembly, which are mainly enriched in pathways related to sensory systems, immunity, etc. This may also be the reason why *G. japonicus* possesses features such as agility, damage repair, and tail regeneration. An important part of these gene sets is T2R, which we then analyzed in more detail.

It has long been shown that T2Rs, genes related to bitter taste perception, are widespread in vertebrates, varies widely among species and that lineage-specific gene duplication events have been detected in bony fishes, frogs, mammals and lizards (Dong et al. 2009). A recent comprehensive analysis by Zhong et al. (Zhong et al. 2019) for T2Rs in squamates has demonstrated that T2Rs are extensively duplicated and expanded in several gecko and lizard lineages, and our location distribution and phylogenetic analyses suggest that this expansion is likely to result from tandem duplication in multiple species. We found that most of the branches under positive selection were located within clades related to *G. japonicus* and *A. carolinensis*, which is concordant with their extensive duplication and expansion, and all these results suggest that T2Rs are important for the adaptive evolution in *G. japonicus*, or insectivorous lizards and geckos. Zhong et al. (Zhong et al. 2019) also demonstrated that the number of T2Rs in different squamates species is closely related to their dietary habits, which is consistent with previous analyses of correlations between vertebrate T2R numbers and dietary habits (Li and Zhang 2014). This is why insectivorous lizards and geckos have such a high number of T2R genes, which are needed for these species to recognize poisonous substances from insects by the perception of bitter taste. The selection pressure on the extracellular, transmembrane and intracellular regions of T2Rs was found to be different in some primates and mammals, and without exception, the extracellular region was favored by positive selection pressure, while the transmembrane region and intracellular region was more favored by purifying selection (Shi et al. 2003; Strotmann et al. 2011; Dong et al. 2021). In sauropsids, we analyzed the selection pressure in extracellular, transmembrane and intracellular regions by the paired YN00 model, which still showed the same pattern as above. The formation of this pattern may be related to the fact that extracellular regions are predicted to be involved in ligand binding, such as tastants (Adler et al. 2000; Gilbertson et al. 2000), and positive selection exerted on this region may facilitate adaptation to different ligand binding. The important changes in the ER region that could affect the ability of species to recognize bitter substances or toxic substances.

There were obviously many birth and death processes in CYPs of sauropsids. Many identified CYPs in sauropsids are not orthologous to that in human, and they may arise through a series of birth and death processes. The MIPhy instability score and the number of gene duplications and losses suggest that birth-death processes apparently occur more frequently in D-type CYP genes. The difference in mean ω values was not significant in D-type CYP genes versus B-type CYP genes, but it correlated with MIPhy scores. We found a close number of D-type and B-type CYP genes under positive selection using the branch-site model, while positive selection branches were found more often among D-type CYP genes using the aBSREL model. We expected to find a correlation between their number and diet in D-type CYPs, as found in T2Rs discussed in the previous section, nevertheless only a weak correlation was found in CYP1s. These results suggest that the interaction between natural selection and birth and death processes of CYPs in sauropsids is complex and interactive, but birth and death processes cannot be fully attributed to the effect of selection pressure, as previously found in CYPs from other groups (Dermauw et al. 2020).

In the present study, 66 CBPs were identified in *G. japonicus*, which are located in the EDC region of chromosome 1 and differed in length. This is in agreement with previous work finding CBPs clustered in the same scaffold of the *G. japonicus* genome (Liu et al. 2015a). In this regard, the formation of specific epidermis in the reptiles are composed of the CBPs coded in the certain region of the chromosome. Among the 66 CBPs, Pro, Cys, Gly and Ser are the most abundant amino acids, and the abundance of Gly and Cys have significant effects on the flexibility and adhesiveness of gecko setae (Alibardi 2013). This results suggest that the CBPs in *G. japonicus* play important roles in the formation of setae in toepads. Multiple sequence alignments revealed the existence of a conserved core box region in the CBPs of *G. japonicus*, which is well conserved in sauropsids and determines the polymerization of these CBP proteins into filaments (Alibardi et al. 2009). Based on the phylogeny, motif, domain and gene structure, we found that the CBP genes family in *G. japonicus* can be divided into three categories in general. Whether these categories correspond to different functions remains to be further studied. Among the studied reptile species, the phylogenetic relationship analysis indicated their CBP genes were not clustered exactly according to species relationships, which suggests that CBP genes are duplicated independently across multiple reptile lineages, leading to variability in the epidermis of reptiles. Therefore, the CBPs in different sauropsids make contributions to the formation of their epidermis respectively. The synteny analysis revealed a high similarity of CBPs in *G. japonicus* and lizards, which is consistent with their morphological similarity, and species relationship.

The results of TRP genes identification in sauropsids are partly consistent with that of TRP analysis in recent study on tuatara (Gemmell et al. 2020), and a certain degree of difference is acceptable considering the differences in methods, but we believe that there are clearly some errors in their results, for example, they did not identify TRPA1 in *A. mississippiensis* and *S. punctatus*, which is a very conserved gene. In our study, it was also identified in each species. In phylogeny, with the exception of TRPV5 and TRPV6, which are thought to be generated by independent duplication in the ancestors of the respective groups of mammals, sauropsids, amphibians, and chondrichthyes (Flores-Aldama et al. 2020), and a low quality TRPV4 sequence of *S. punctatus*, each member of the other TRP families are well clustered into a monophyletic group. In *G. japonicus*, TRPV5, TRPM5, and TRPM8 were under relaxed purifying selection, while TRPM3 and PKD2L2 were under positive selection and MCOLN1 was under intensified purifying selection. TRPV5 is a channel that exhibits high selectivity for calcium ions, mediates calcium ion influx into cells, and is responsive to activation and expression of hormones such as parathyroid hormone, estrogen and testosterone (Na and Peng 2014), and the relaxation of purified selection suggests that possibly the regulation of expression of these hormones, undergoes some adaptive changes among *G. japonicus*. MCOLN1 (TRPML1) has been revealed to be associated with lysosomal function in mammals, and its abnormalities can cause disorders of lysosomal channels and function, leading to lysosomal dysfunction and lysosomal storage diseases, and may also be responsible for the pathogenesis of metabolic and common neurodegenerative diseases (Xu and Ren 2015). The importance of lysosomal function in *G. japonicus* could make it more favored for purification selection. PKD2L2 plays a important role in reproduction and is an essential gene for spermatogenesis (Chen et al. 2008; Dolebo et al. 2019), which suggests that the reproductive system of *G. japonicus* have undergone adaptive evolution. PKDREJ and PKD2L2, two reproduction-related genes, have higher ω values relative to other TRP genes, which is consistent with previous studies finding that reproduction-related proteins have higher evolutionary rates (Swanson et al. 2001; Swanson and Vacquier 2002; Clark et al. 2006; Walters and Harrison 2010; Walters and Harrison 2011; Wilburn and Swanson 2016; Twort et al. 2017). TRPM3, TRPM5 and TRPM8 are thermo-sensitive TRP genes (Wang and Siemens 2015), which suggests that their genomic changes may be related to the evolution of thermoregulation in *G. japonicus*. Compared to other reptile species, *G. japonicus* is a relatively widely distributed species, and there are various reasons that can allow a species to adapt and survive in a wide range, and some of the thermosensitive TRPs under relaxed purifying selection or positive selection in *G. japonicus*, implying that one of the important factors is the ability to adapt to temperature changes. In fact, in 2020, several researchers predicted the present and future distribution of *G. japonicus* and found that temperature seasonality was an important predictor of its distribution (Kim et al. 2020), also indicating the process of adaptive evolution to temperature in *G. japonicus*. These thermosensitive TRPs which are under relaxed purifying selection or positive selection may have corresponding changes in the range and magnitude of activation temperature due to the wide variation of environmental temperature. Our results provide evidence for the temperature adaptation of *G. japonicus* and reveal a possible reason for its wide distribution.

## Conclusions

In this study, we assembled and annotated a high quality *G. japonicus* genome with 19 chromosomes based on Illumina sequencing, PacBio SMRT sequencing and Hi-C technology. To our knowledge, our study represents the first chromosomal level assembly of the gecko genome. Comparative genomic analyses revealed the phylogenetic position of sauropsids, important functional gene sets that are mostly associated with immune and sensory systems and enriched to related pathways, chromosomal rearrangement events and chromosome evolution in *G. japonicus*, implying important molecular mechanisms for tail regeneration, damage repair, and sensitive senses in *G. japonicus*. The evolutionary trajectories of bitter taste receptor type 2 (T2Rs), detox and biosynthetic cytochrome P450 (CYPs), epidermis formation corneous beta proteins (CBPs) and thermoregulatory transient receptor potential channels (TRPs) in *G. japonicus* and other representative sauropsids are identified. These results indicated that T2Rs expanded in different lineages by tandem gene duplication. The expansion and independent duplication events of the T2Rs and positively selected branches were predominantly present in insectivorous species, revealing an important relevance of the T2Rs to the clearance of bitter toxins in gecko. D-type genes in CYP family of *G. japonicas* have frequent duplication and loss events, which suggests that they undergo more birth and death processes compared to B-type CYP genes. Pro, Cys, Gly and Ser are the most abundant amino acids in 66 CBPs of *G. japonicas*, the abundance of Gly and Cys in CBPs implying significant effects on the flexibility and setae adhesiveness of *G. japonicas*. Some thermosensitive TRPs under relaxed purifying selection or positive selection in *G. japonicus*, implying that one of the important factors improve the ability to adapt to climate change. In summary, we assembled a high-quality chromosome-level genome of *G. japonicus* and performed a comprehensive comparative genomics and evolutionary genomics analyses of *G. japonicus*, and revealed the regional characterization and evolutionary contribution of EBRs, the important relevance of T2Rs and CYPs in the detoxification mechanism of insectivorous species and the possible role of CBPs in phenotypic adaptation evolution of toepads and TRPs in temperature perception and regional expansion in gekkotans.

## Methods

### Sampling and sequencing

The geckos were acquired from various localities in Nanjing (32°03′N, 118°45′E), eastern China and transported to our laboratory at Nanjing Normal University in 500 mm × 300 mm × 250 mm (length × width × height) plastic cages. They were fed with mealworms and given water regularly during the whole experiment. In the present work, the affidavit of Approval of Animal Ethical and Welfare were approved by the Institutional Animal Care and Use Committee of Nanjing Normal University [SYXK (Jiangsu) 2020–0047 and IACUC-20220258]. One of the healthy adult female gecko was chosen to prepare the muscle and visceral tissue. In order to minimize the degradation of DNA and RNA, the gecko was transported with dry ice and then preserved in the refrigerator at −80°C. Then its muscle and visceral tissue were rapidly collected in the clean bench. After weighing, 2.06 g of muscle tissue and 0.90 g of visceral tissue were put into two 1.5 mL centrifuge tubes respectively. The labels were marked on the tubes and immediately put in the foam box filled with liquid nitrogen for preservation. The box was sent to Wuhan Frasergen Bioinformatics Co., Ltd. for further DNA extraction, RNA extraction, Hi-C cell crosslinking experiment and Illumina Hiseq sequencing, PacBio SMRT sequencing and Hi-C analysis.

### Genome assembly and annotation

Genome size was estimated by K-mer (17-mer) analysis using reads generated by Illumina Hiseq sequencing according to Lander-waterman theory. The preliminary genome assembly were first obtained using MECAT2 (Xiao et al. 2017) based on reads generated by PacBio SMRT sequencing, GCpp version 1.9.0 (https://github.com/PacificBiosciences/gcpp) was used to polish the assembly using subreads, then Pilon version 1.22 (Walker et al. 2014) was used to correct the errors based on Illumina data, and finally Hi-C analysis was performed using juicer version 1.6.2 (Durand et al. 2016) to obtain a chromosome-level genome assembly.

Homology prediction of repetitive elements using RepeatMasker and RepeatProteinMask version 4.0.9 (Price et al. 2005; Tarailo-Graovac and Chen 2009) based on RepBase (Jurka et al. 2005), repetitive elements were predicted based on their own sequence comparison and de novo prediction using RepeatModeler version 1.0.11 (Smit and Hubley 2008), and using LTR-FINDER version 1.0.5 (Xu and Wang 2007) for de novo prediction based on repeated sequence features. In addition, the de novo prediction approach also used TRF version 4.09 (Benson 1999) to find tandem repeat sequences in the genome. These results were integrated to obtain the final repeated sequence annotation.

Using MAKER pipeline version 3.01.02 (Cantarel et al. 2008) based on exonerate version 2.2.0 (Slater and Birney 2005) with homologous proteins of *Podarcis muralis*, *Anolis carolinensis*, *Pogona vitticeps*, *Lacerta agilis* and *Gekko japonicus* (previous version), third-generation full-length transcriptome data, second-generation transcriptome data, and the ab initio prediction software Augustus version 3.3 (Stanke and Waack 2003), Genscan version 1.0 (Burge and Karlin 1997) to obtain a non-redundant complete gene set. Benchmarking Universal Single-Copy Orthologs (BUSCO) version 2.0 (Simão et al. 2015) was used to assess genome assembly and gene annotation. The proteins in the gene set were functionally annotated based on protein databases KEGG (Kanehisa and Goto 2000), InterPro (Mitchell et al. 2019), GO (Consortium 2004), NR (Marchler-Bauer et al. 2015), SwissProt and TrEMBL (Boeckmann et al. 2003). Non-coding RNAs were annotated based on tRNAscan-se version 1.3.1 (Lowe and Eddy 1997), BLASTN in BLAST+ version 2.6.0 (Camacho et al. 2009) and Rfam version 14.1 (Griffiths-Jones et al. 2005). A search for 1:1 orthologous genes between two *G. japonicus* genome assemblies was performed using OrthoFinder version 2.5.1 (Emms and Kelly 2015). Synteny analysis was performed using BLASTP (e-value = 1e-5) and MCScanX (Wang et al. 2012), other genomic features were calculated using BEDTools version 2.29.2 (Quinlan 2014) based on our annotations, and finally syntenic regions and genomic features were plotted using Circos version 0.69 (Krzywinski et al. 2009).

### Synteny analysis and identification of HSBs and EBRs

OrthoFinder version 2.5.1 (Emms and Kelly 2015) was used to identify the orthologous genes of *G. japonicus* with *A. carolinensis* and *G. japonicus* with *G. gallus*. The chromosome sequences and orthologous genes of each of these three species were used as input to Symap version 5.0.6 (Soderlund et al. 2006) to identify HSBs of *G. japonicus* with *A. carolinensis* and *G. japonicus* with *G. gallus*, and the syntenic regions were visualized with Circos version 0.69 (Krzywinski et al. 2009). The regions between adjacent HSBs were the EBRs. GO enrichment was performed with clusterProfiler version 4.0 (Wu et al. 2021) in R version 4.1.0. The wilcoxon rank sum test was performed in R version 4.1.0 to test whether the gene density, GC content and repeat content of the EBR of each chromosome were significantly different from those of the whole chromosome. Linear regression analysis was performed in Python version 3.8.1 using the module statsmodel and visualized with the seaborn module. A t-test with unequal variances was used to test whether each repetitive element in EBRs was significantly higher than the same repetitive element in the rest of the genome, which was calculated based on 10-kbp windows, performed in R version 4.1.0.

### Phylogenetic and gene family analysis

A total of 22 species genomes were selected for gene family and phylogenetic analyses, including 4 species of snakes (*Ophiophagus Hannah*, *Protobothrops mucrosquamatus*, *Python bivittatus* and *Pantherophis guttatus*), and 5 species of lizards (*Anolis carolinensis*, *Podarcis muralis*, *Pogona vitticeps*, *Gekko japonicus* (this study), *Gekko japonicus* (previous version)), 3 species of birds (*Gallus Gallus*, *Meleagris gallopavo*, *Taeniopygia guttata*), 4 species of crocodiles (*Alligator misissippiensis*, *Alligator sinensis*, *Crocodylus porosus*, and *Gavialis gangeticus*), three species of turtles (*Chelonia mydas*, *Chrysemys picta bellii*, and *Perodiscus sinensis*), two species of mammals (*Homo sapiens* and *Mus musculus*) and an amphibian (*Xenopus tropicalis*).

Gene family clustering is an important step in phylogenetic analysis. Protein sequences of various species were clustered based on sequence similarity by OrthoMcl version 1.4 (e-value of BLASTP is 1e-5 and alignments with identity < 30% or coverage < 50% were dropped) (Li et al. 2003). The single-copy orthologous genes were filtered, and only the genes with amino acid length ≥100 were retained. Muscle version 3.8.31 (Edgar 2004) was used to perform multiple sequence alignment for each single-copy orthologous gene set and the alignments were concatenated and converted into a super-gene alignment in Phylip format.

RAXML version 8 (Stamatakis 2014) was used to construct the phylogenetic tree based on the maximum likelihood method. Using the phylogenetic tree constructed, combined with TimeTree (Kumar et al. 2017) and literature to obtain time correction points, the MCMCTree program in PAML version 4.7 (Yang 2007) and R8s version 1.50 (Sanderson 2003) was used to estimate the divergence time. CAFE version 4.2.1 (De Bie et al. 2006) was used to estimate the expansion and contraction events of gene families by employing a random birth and death model to find gene gain and loss across the phylogenetic tree. According to the results of gene family clustering, for each single-copy orthologous gene set, the branch-site model in PAML version 4.7 (Yang 2007) was used to detect whether genes are under positive selection in *G. japonicus*. This model is very useful for testing positive selection on specific branches,which assumes four classes of sites, the first class of sites 0 is assumed to have 0 < ω_0_ < 1 in all branches, the second class of sites 1 is assumed to have ω_1_ = 1 in all branches, the third class of sites 2a is assumed to have ω_2a_ = ω_2b_ ≥ 1 in the foreground and 0 < ω_2a_ = ω_0_ < 1 in the background, and the fourth class of sites 2b is assumed to have ω_2b_ = ω_2a_ ≥ 1 in the foreground and ω_2b_ = ω_1_ = 1 in the background. It only allows positive selections in the foreground, but may result in false positives when positive selection is present in the background (Schott et al. 2014; Schott et al. 2018). Go/KEGG enrichment analysis was performed by Fisher’s exact test for the expanded and contracted gene families, specific gene families and positively selected genes in *G. japonicus*, respectively. P-value was then corrected based on FDR method to obtain q-value, and results less than 0.05 were retained.

### Analysis of T2Rs

Seven sauropsids, including *Anolis carolinensis*, *Alligator sinensis*, *Gallus gallus*, *Gekko japonicus*, *Python bivittatus*, *Pelodiscus sinensis*, and *Sphenodon punctatus*, across all extant taxa, as representative small data sets for gene identification, to compare and complement previous gene identification results, and for evolutionary analysis.

The T2R protein sequences of western clawed frog, human, mouse and chicken were downloaded from NCBI and used them as reference sequences to identify the T2R sequences of other species using GeMoMa version 1.7.1 (Keilwagen et al. 2016) by TBLASTN method with the threshold set as 1e-10. Next, the transmembrane regions of the candidate sequences were predicted using TMHMM version 2.0 (Krogh et al. 2001), and only sequences with transmembrane number greater than or equal to 7 were retained. To ensure the high quality and structural integrity of the sequences, we also removed sequences with codon number less than or equal to 270, and finally obtained a high quality T2R gene set containing only intact genes, which all begin with a start codon and end with a stop codon, and then named the genes with species name acronyms and numbers for subsequent analysis. We used MACSE version 2.05 (Ranwez et al. 2018) to align the T2R coding DNA sequences, and then manually removed low quality regions in Mega-X version 10.0.5 (Kumar et al. 2018). The phylogenetic tree of the T2R proteins alignment for seven species was constructed using RAxML-NG v. 0.9.0 (Kozlov et al. 2019) based on the best model JTT+I+G4+F selected by ModelTest-NG version 0.1.7 (Darriba et al. 2020). Subtree of *G. japonicus* was extract by ETE 3 (Huerta-Cepas et al. 2016). All trees were edited in Adobe Illustrator CC 2019, iTOL (Letunic and Bork 2021) and R package ggtree (Yu et al. 2017). M3 vs. M0, M2a vs. M1a, and M8 vs. M7/M8a in PAML version 4.7 (Yang 2007) and FEL (Kosakovsky Pond and Frost 2005) and FUBAR (Murrell et al. 2013) in HyPhy version 2.5.2 (Kosakovsky Pond et al. 2020) were performed to identify positively selected sites in T2R genes. The aBSREL (Smith et al. 2015) in HyPhy version 2.5.2 (Kosakovsky Pond et al. 2020) was used to find some branches in the T2R tree of 7 species and in the T2R tree of *G. japonicus* that were under positive selection. Multiple sequence alignments of the T2R gene were divided into ER, TM and IR regions with reference to a chicken T2R protein (XP_004938201.1 or Q2AB83) in UniProt (Consortium 2015). YN00 in PAML version 4.7 (Yang 2007) was used to calculate pairwise ω values in ER, TM and IR regions. All statistical tests were performed in R version 4.1.0.

### Analysis of CYPs

Functional CYP gene sequences of human were obtained from the Cytochrome P450 Homepage (Nelson 2009), then we used GeMoMa version 1.7.1 (Keilwagen et al. 2016) with TBLASTN method to search the genomes of 13 sauropsids including two crocodiles (*Alligator misissippiensis* and *Alligator sinensis*), two lizards (*Anolis carolinensis* and *Pogona vitticeps*), two geckos (*Gekko japonicus* (this study) and *Paroedura picta*), two snakes (*Vipera berus* and *Ophiophagus hannah*), three turtles (*Chelonia mydas*, *Chrysemys picta bellii*, and *Perodiscus sinensis*), tuatara (*Sphenodon punctatus*) and one bird (*Gallus gallus*), using the CYP protein sequences of human as references. We filtered out protein sequences with lengths less than 350, then aligned them with *Drosophila melanogaster* CYP301a1 using MAFFT version 7.475 (Katoh and Standley 2013) (strategy L-INS-i). We used trimAl version 1.4.rev15 (Capella-Gutierrez et al. 2009) (gappyout) to trim low quality regions in the protein alignmnet. RAxML-NG v. 0.9.0 (Kozlov et al. 2019) was used to build ML tree using the best model JTT+G4 found by ModelTest-NG version 0.1.7 (Darriba et al. 2020) and 1000 repeated samples to test the confidence of each branch, and *D. melanogaster* CYP301a1 as root of the tree.

MIPhy was used to identify clades and assess their instability scores, and then PAML version 4.7 (Yang 2007) and HyPhy version 2.5.2 (Kosakovsky Pond et al. 2020) were used to perform selection pressure analysis for each clade. M0 model in PAML was used to calculate average ω for each clade, and M2a vs. M1a and M8 vs. M7 in PAML was used to test to test if there are positively selected sites in each clade. The branch-site model in PAML was used to test whether branches of *G. japonicus* were under positive selection, and the aBSREL model in HyPhy was used to find positively selected branches in each clade. NOTUNG version 3.0.26 (Chen et al. 2000) was used to identify gene duplication and loss events in sauropsids CYPs, PICs was calculated in R package ape (Paradis and Schliep 2019) and all other statistical tests were run in R version 4.1.0.

### Analysis of CBPs

The identified CBP gene family sequences of *Gekko gecko* (Hallahan et al. 2009) and *A. carolinensis* (Dalla Valle et al. 2010) were used as reference to search the third-generation genome sequences of *G. japonicus* was used as database to extract the CBP gene family of *G. japonicus* by BLASTP, setting e-value as 10-5. Then, the genomes and annotated data of other selected species were download, they were *Paroedura picta*, *Anolis carolinensis*, *Podarcis muralis*, *Zootoca vivipara*, *Lacerta agilis*, *Pogona vitticeps*, *Python bivittatus*, *Ophiophagus hannah*, *Thamnophis elegans*, *Pelodiscus sinensis*, *Chrysemys picta*, *Chelonia mydas*, *Alligator sinensis*, *Alligator mississippiensis*, *Crocodylus porosus*, *Xenopus tropicalis*, *Mus musculus*, *Eublepharis macularius* and *Shinisaurus crocodilurus*. The same method was performed to obtain the CBP gene family in each species mentioned above. 66 amino acid sequences of CBP gene family members of *G. japonicus* were obtained.

The physicochemical properties and components of CBP amino acid sequences obtained from *G. japonicus* were analyzed. With online website ProtParam (https://web.expasy.org/protparam/) to calculate the basic information of CBP amino acid sequences such as the molecular weight (MW) and isoelectric point (PI). The amino acid composition ratio of each sequence was calculated using local software Mega-X version 10.0.5 (Kumar et al. 2018). The motif of each sequence was analyzed by MEME online page (http://meme-suite.org/). Then the conservative domain characteristics of the sequences were analyzed by NCBI online tool (https://www.ncbi.nlm.nih.gov/cdd/). The local software TBtools version 1.074 (Chen et al. 2020) was used to visually analyze the gene structure and the distribution of motif and domain in each sequence of the acquired gene family members. R packages ggplot2 and gggenes were used to plot and analyze the position, arrangement and orientation of gene family members in the chromosomes.

Phylogenetic analysis was performed among the CBP gene family members of *G. japonicus*. MAFFT version 7.475 (Katoh and Standley 2013) (strategy L-INS-i) was used for multi-sequence alignment. ModelTest-NG version 0.1.7 (Darriba et al. 2020) was used to calculate the optimal model of constructing Maximum Likelihood (ML) tree. RAxML-NG v. 0.9.0(Kozlov et al. 2019) was used to build ML tree based on the best model WAG+G4+F, and 1000 repeated samples were used to test the confidence of each branch. ML trees were constructed for all selected reptile species in the same way to explore the evolutionary origin of the CBP gene family of *G. japonicus*.

In order to compare the number and distribution of CBPs among selected species, regional synteny analysis was performed on these sequences. Sequences and gene annotation files inside the flanking genes of the CBP sequences were intersected by BEDTools version 2.29.2 (Quinlan 2014). Synteny analysis of the CBP gene families between two species was performed by MCScanX (Wang et al. 2012). TimeTree (Kumar et al. 2017) was used to construct the time tree between the selected species, which was used as the basis for drawing the sequence of gene family arrangement map of the species.

### Analysis of TRPs

TRP genes of representative sauropsids, two mammals and one amphibians were obtained from NCBI, including one frog (*Xenopus tropicalis*), human (*Homo sapiens*), mouse (*Mus musculus*), two birds (*Gallus gallus* and *Taeniopygia guttata*), one lizard (*Anolis carolinensis*), one gecko (*Gekko japonicus*, previous version), one snake (*Python bivittatus*), three turtles (*Chelonia mydas*, *Chrysemys picta* and *Pelodiscus sinensis*) and four alligators (*Alligator mississippiensis*, *Alligator sinensis*, *Crocodylus porosus* and *Gavialis gangeticus*). Then we used GeMoMa version 1.7.1 (Keilwagen et al. 2016) with TBLASTN method to search the genomes of two geckos (*Gekko japonicus* (this study) and *Paroedura picta*), two snakes (*Vipera berus* and *Ophiophagus hannah*) and tuatara (*Sphenodon punctatus*) using the TRP protein sequences of *Xenopus tropicalis*, *Homo sapiens*, *Gallus gallus*, *Anolis carolinensis* and *Gekko japonicus* (previsou version) as references. TMHMM version 2.0 (Krogh et al. 2001) was used to predict TM topology, sequences which did not have at least 3 predicted TM segments were removed (TRP commonly has 6 transmembrane regions, here we chose a lower threshold to retain some partial and intact sequences). The TRP protein sequences of 19 species and the VDAC protein sequences of *Xenopus tropicalis* as an outgroup were aligned using MAFFT version 7.475 (Katoh and Standley 2013) (strategy L-INS-i). The aligned sequences were trimmed using the trimAl version 1.4.rev15 (Capella-Gutierrez et al. 2009) (gappyout). The maximum likelihood tree was constructed using RAxML-NG version 0.9.0 (Kozlov et al. 2019) (1,000 bootstraps) with the best-fit model JTT+G4+F selected by ModelTest-NG version 0.1.7 (Darriba et al. 2020), and rooted with the VDAC protein sequences of *Xenopus tropicalis*. The predicted genes were named by their phylogenetic relationships and BLASTP results. We visualized and edited our tree in Adobe Illustrator CC 2019, iTOL (Letunic and Bork 2021) and R package ggtree (Yu et al. 2017). Single TRP subfamily members were aligned using the Prank_+F_ (Löytynoja and Goldman 2008) algorithm in GUIDANCE2 (Sela et al. 2015), and unreliable codons (with scores < 0.9) were replaced with “NNN” based on the residue filtering principle. Four pairs of site models M3 vs. M0, M2a vs. M1a, and M8 vs. M7/M8a in PAML version 4.7 (Yang 2007) were performed to identify positively selected sites and calculate average ω values for each TRP gene. CmC and CmD model in PAML version 4.7 (Yang 2007) are well suited to test for divergent selection, and we used them to test for shifts in selection pressures. CmC assumes that throughout the phylogeny one class of sites has a ω value greater than 0 and less than 1, another class of sites has a ω value equal to 1, and in two or more partitions one class of sites is free to evolve differently with unconstrained ω values. CmD is similar to CmC, but all three classes of sites are unconstrained. We also used branch-site model and branch model in in PAML version 4.7 (Yang 2007) to test for positive selection and relaxed selection. The branch model is very useful to test relaxed selection, it can calculate average ω values between different partitions. All models were run at least three times with different initial values of κ and ω (except for some null models with fixed values) to ensure the robustness of the results. FDR control was performed with R package qvalue based on the method of Storey et al (Storey 2002). We also performed Bonferroni correction with the similar results in R version 4.1.0. YN00 in PAML version 4.7 (Yang 2007) was used to calculate the pairwise ω values of 1:1 orthologous TRP genes between *Gekko japonicus* and *Anolis carolinensis*. All other statistical tests were run in R version 4.1.0.

## Data access

All raw sequencing reads have been deposited in the NCBI Sequence Read Archive under project PRJNA899667. Predicted gene sequences, sequence alignments and phylogenetic trees can be downloaded from https://doi.org/10.5281/zenodo.7211720. The chromosome-scale genome assembly and annotation files are available from https://doi.org/10.5281/zenodo.7394737.

## Acknowledgments

This work was supported by grants from the National Natural Science Foundation of China (Grant No. 31672269 to JY), Postgraduate Research & Practice Innovation Program of Jiangsu Province (KYCX22_1614) awarded to ZYC, the Priority Academic Program Development of Jiangsu Higher Education Institutions (PAPD).

## Author contributions

PL and JY conceived and designed the experiments. YWW and YXY performed the experiments. YWW, YXY, CL, ZYC, YC, CCH, YFQ and HL analyzed data and prepared all of figures and tables. YWW, YXY, JY, and PL wrote and revised the manuscript. All authors discussed the results and implications and commented on the manuscript.

## Author details

Yinwei Wang is a herpetologist at Nanjing Normal University, he is interested in ecology and evolutionary biology of reptiles. Youxia Yue is a herpetologist at Nanjing Normal University, she is interested in molecular ecology and evolutionary biology of reptiles. Chao Li is a herpetologist at Nanjing Normal University, she is interested in molecular evolution, ecology and evolutionary biology of reptiles. Zhiyi Chen a herpetologist at Nanjing Normal University, she is interested in molecular evolution, ecology and evolutionary biology of reptiles. Yao Cai is a herpetologist at Nanjing Xiaozhuang University, he is interested in physiological ecology and evolutionary genetics, and biodiversity of reptiles. Chaochao Hu is a herpetologist and ornithologist at Nanjing Normal University, he is interested in diversity, ecology and evolution of reptiles and birds. Yanfu Qu is a herpetologist at Nanjing Normal University, he is interested in intestinal microorganism, physiology, ecology and evolution of reptiles. Hong Li is a herpetologist at Nanjing Normal University, he is interested in physiology, ecology and evolution of reptiles. Kaiya Zhou is a marine mammals zoologist and herpetologist at Nanjing Normal University, he is especially interested in the conservation biology, habitats, and regions, niche evolution, and molecular phylogenetics and evolution. Jie Yan a herpetologist at Nanjing Normal University, she is interested in molecular evolution, phylogeny and biogeography of amphibians and reptiles. Peng Li is a herpetologist at Nanjing Normal University, he is interested in molecular evolutionary mechanisms of phenotypic plasticity, comparative genomic and evolutionary genetics, phylogeny and biogeography, and biodiversity and distribution patterns of reptiles.

## Supplemental information

**Supplemental_Figure_S1.docx:** Supplemental **Figures S1-S20**.

Figure S1. Evaluation of *Gekko japonicus* genome size by k-mer analysis.

Figure S2. Hi-C contact heatmap for *Gekko japonicus*.

Figure S3. Box plot and violin plot of 1:1 orthologous gene length for two different genome assembly versions.

Figure S4. Box plot and violin plot of 1:1 orthologous protein length for two different genome assembly versions.

Figure S5. Linear regression analysis of repetitive sequence content and gene density in EBRs.

Figure S6. GO enrichment analysis of the Gekko japonicus-specific EBRs.

Figure S7. GO enrichment analysis of the *Gekko japonicus*-specific expanded gene families.

Figure S8. KEGG enrichment analysis of the *Gekko japonicus*-specific expanded gene families.

Figure S9. GO enrichment analysis of genes under positive selection in *Gekko japonicus*.

Figure S10. KEGG enrichment analysis of genes under positive selection in *Gekko japonicus*.

Figure S11. GO enrichment analysis of the *Gekko japonicus*-specific gene families.

Figure S12. Instability scores of CYP450 clades.

Figure S13. Box plot of the number of each family for sauropsids CYP450 genes.

Figure S14. Relationship between the number of CYPs and Diet.

Figure S15. Amino acid composition of 66 CBPs in *Gecko japonicus*.

Figure S16. The “core box” motif of 66 CBPs alignment of *Gecko japonicus*.

Figure S17. The motif distribution of 66 CBPs in *Gecko japonicus*.

Figure S18. Phylogenetic tree, motif distribution, domain distribution and gene structure distribution of 66 CBPs of *Gekko japonicus*.

Figure S19. Location of the CBP gene family in 20 species.

Figure S20. Pairwise covariance analysis of the CBP gene family of 19 species with that of the *Gekko japonicus*.

**Supplementary_Table_S1.xlsx**: Supplemental **Tables S1-S63**.

Table S1. Lengths of the 19 assembled chromosomes in *Gekko japonicus*.

Table S2. Statistics for scaffolds and contigs in the genome assembly of *Gekko japonicus*.

Table S3. Assessment of genome assembly and annotation using BUSCO.

Table S4. Statistics of repeated sequences results based on four methods.

Table S5. Classification statistics of transposable elements (TEs).

Table S6. Results of gene prediction based on multiple software and methods.

Table S7. Summary of results for non-coding RNA annotation.

Table S8. Comparison of our protein-coding genes annotation with previous version.

Table S9. Summary of results for functional annotations.

Table S10. Position information of HSBs between *Gekko japonicus* and *Anolis caroliensis*.

Table S11. Position information of HSBs between *Gekko japonicus* and *Gallus gallus*.

Table S12. EBRs in *Gekko japonicus* identified by HSBs between *G. japonicus* and *A. carolinensis*.

Table S13. EBRs in *Gekko japonicus* identified by HSBs between *G. japonicus* and *G. gallus*.

Table S14. The merged EBRs obtained by combining the EBR regions of the two alignments.

Table S15. Comparison of gene density, GC content and repeat content across chromosomes between EBRs and genome.

Table S16. Densities of repetitive element families per 10-kbp in specific EBRs of *Gekko japonicus*.

Table S17. Accessions of T2R protein sequences used to query.

Table S18. Results of PAML site model analyses of T2Rs for seven sauropsids.

Table S19. Positively selected sites of T2Rs in seven sauropsids identified by FEL model.

Table S20. Positively selected sites of T2Rs in seven sauropsids identified by FUBAR model.

Table S21. Results of PAML site model analyses of T2Rs for *Gekko japonicus*.

Table S22. Positively selected sites of T2Rs in *Gekko japonicus* identified by FEL model.

Table S23. Positively selected sites of T2Rs in *Gekko japonicus* identified by FUBAR model.

Table S24.Gene repertoire of CYP450 in 13 sauropsids.

Table S25. Duplications and losses of CYP450 genes in 13 sauropsids.

Table S26. Site-model analysis for sauropsids CYPs.

Table S27. Positively selected branches found in sauropsids CYPs using aBSREL Model.

Table S28. Branch-site model analysis for sauropsids CYPs.

Table S29. Location and basic information of 66 CBPs sequences of *Gecko japonicus*.

Table S30. Accessions of the TRPs researched in this study.

Table S31. PAML analyses for TRPA1.

Table S32. PAML analyses for TRPC1.

Table S33. PAML analyses for TRPC2.

Table S34. PAML analyses for TRPC3.

Table S35. PAML analyses for TRPC4.

Table S36. PAML analyses for TRPC5.

Table S37. PAML analyses for TRPC6.

Table S38. PAML analyses for TRPC7.

Table S39. PAML analyses for TRPM1.

Table S40. PAML analyses for TRPM2.

Table S41. PAML analyses for TRPM3.

Table S42. PAML analyses for TRPM4.

Table S43. PAML analyses for TRPM5.

Table S44. PAML analyses for TRPM6.

Table S45. PAML analyses for TRPM7.

Table S46. PAML analyses for TRPM8.

Table S47. PAML analyses for TRPV1.

Table S48. PAML analyses for TRPV2.

Table S49. PAML analyses for TRPV3.

Table S50. PAML analyses for TRPV4.

Table S51. PAML analyses for TRPV5.

Table S52. PAML analyses for TRPV6.

Table S53. PAML analyses for MCOLN1 (TRPML1).

Table S54. PAML analyses for MCOLN2 (TRPML2).

Table S55. PAML analyses for MCOLN3 (TRPML3).

Table S56. PAML analyses for PKD1.

Table S57. PAML analyses for PKD1L2.

Table S58. PAML analyses for PKD2.

Table S59. PAML analyses for PKD2L1.

Table S60. PAML analyses for PKD2L2.

Table S61. PAML analyses for PKDREJ.

Table S62. P-value of LRTs in PAML analysis.

Table S63. Q-value of LRTs in PAML analysis.

## Notes

### Competing Interest Statement

The authors have declared no competing interest.

